# A mechano-osmotic feedback couples cell volume to the rate of cell deformation

**DOI:** 10.1101/2021.06.08.447538

**Authors:** Larisa Venkova, Amit Singh Vishen, Sergio Lembo, Nishit Srivastava, Baptiste Duchamp, Artur Ruppel, Stéphane Vassilopoulos, Alexandre Deslys, Juan Manuel Garcia Arcos, Alba Diz-Muñoz, Martial Balland, Jean-François Joanny, Damien Cuvelier, Pierre Sens, Matthieu Piel

## Abstract

Mechanics has been a central focus of physical biology in the past decade. In comparison, the osmotic and electric properties of cells are less understood. Here we show that a parameter central to both the physics and the physiology of the cell, its volume, depends on a mechano-osmotic coupling. We found that cells change their volume depending on the rate at which they change shape, when they spread, migrate or are externally deformed. Cells undergo slow deformation at constant volume, while fast deformation leads to volume loss. We propose a mechano-sensitive pump and leak model to explain this phenomenon. Our model and experiments suggest that volume modulation depends on the state of the actin cortex and the coupling of ion fluxes to membrane tension. This mechano-osmotic coupling defines a membrane tension homeostasis module constantly at work in cells, causing volume fluctuations associated with fast cell shape changes, with potential consequences on cellular physiology.

## Introduction

In recent years, *in vivo* imaging has revealed that, in a variety of physiological and pathological contexts, cells undergo large deformations ^1^, sometimes being squeezed to a tenth of their resting diameter. Migrating cells, in particular fast-moving immune or cancer cells, can deform to a large extent in only a few minutes ^2–4^, for example when they cross an endothelial barrier ^5^. Even faster deformations, below the second timescale, can be observed in circulating cells pushed through small blood capillaries. Altogether, these examples show that large cell deformations are physiological and occur across a large range of timescales. In all these examples, the capacity of cells to adapt and survive large deformations is a key element of physiological and pathological processes. Nevertheless, there is so far very little knowledge on both the physics and the biology of these large cellular deformations.

Large cell shape changes must involve significant changes in volume, surface area, or both. But the number of studies on cell volume modulation upon cell deformation is still very small ^6,7^. It is still not clear whether the material that cells are made of is rather poroelastic ^8^, losing volume when pressed, or behaves like a liquid droplet, extending its surface area at constant volume. Two articles, measuring volume using 3D reconstruction from confocal slices, report that cells that are more spread are smaller in volume ^9,10^, leading to a higher density and potential long-term effects on cell fate ^9^. On the other hand, another article, using volume measured by fluorescence exclusion (FXm), reports no or slightly positive correlation between spreading area and cell volume ^11^, reflecting the fact that as cells grow larger, their spreading area also increases.

Different models have recently been proposed to explain a coupling between cells shape changes and cell volume modulation ^9,10,12,13^. Most of them are based on the same type of scenario: depending on the timescale and extent of the deformation, cell shape changes can stress the cell surface, including the membrane and the actin cortex ^14^. This stress can be relaxed due to cortex turnover, unfolding of membrane reservoirs ^15^ and detachment of the membrane from the cortex with the formation of blebs ^16^. Stress in these structures can also lead to the modulation of ion fluxes ^17^ resulting in cell volume changes. Despite its broad relevance for cell mechanics and cell physiology, the consequences of this type of scenario have not been explored in depth experimentally.

Using FXm to accurately measure volume in live cells ^18–20^, we investigated various experimental contexts in which cells undergo large shape changes: classical osmotic shocks, experimentally imposed deformations (2D confinement) or self-imposed deformations (cell spreading, including in the context of cell migration). We observed that cells modulate their volume at various timescales from milliseconds to minutes. We also found that, during cell spreading, the degree of volume changes depends on both the state of the actomyosin cortex and the rate of deformation. These observations can be explained by an extension of the classical pump and leak model ^21^, including a mechano-osmotic coupling activated upon cell deformation occurring faster than the membrane tension/actin cortex relaxation timescale. We believe that our observations - together with this novel physical model, which we fit and test experimentally - constitute strong evidence for the existence of a mechano-osmotic coupling constantly at work in animal cells and modulating their volume as they deform.

## Results

### Cell volume depends on spreading speed and not on steady-state spreading area

We first asked whether in a population growing and dividing at steady state, cells display a correlation between their spreading area and their volume. We used HeLa cells expressing hGeminin-mCherry, which accumulates in the nucleus during S phase. Cell spreading area was measured using phase contrast and cell volume using FXm ^19^ (**Figure 1A** images). We did not find any strong correlation between spreading area and volume for HeLa and RPE-1 cells, larger cells in volume being also slightly more spread (**Figure 1A** graph and **Figure S1A** and **S1B**). A clearer positive correlation was observed for 3T3 fibroblasts, which were also generally more spread for a given volume (**Figure S1C**). Using the hGeminin cell cycle marker, we observed that S/G2 cells tend to be larger and more spread than G1 cells (**Figure 1A** right), suggesting that the positive correlation is simply due to cell growth, with cells increasing their spreading area as they grow. Using live cell recording of phase, volume and hGeminin, we also considered cells at given windows of time following cell division, to examine the correlation between volume and surface area at a given cell cycle stage and thus independently of cell growth. Considering the same group of cells at various times after mitosis, or after the G1/S transition, we could not observe any correlation between cell volume and spreading area at any given cell cycle stage (**Figure 1A** and **S1D**). Finally, to extend the range of spreading areas considered, we used adhesive micro-patterns with areas smaller than the average spontaneous steady-state spreading area of HeLa cells (**Figure 1B** images). We found that the distribution of volumes did not change when cells were plated on smaller adhesive patterns (**Figure 1B** graph). Overall, these experiments suggest that, as reported before ^11^, there is no strong correlation, at the cell population level, between spreading area and cell volume, independently of the cell cycle stage.

**Figure 1:**
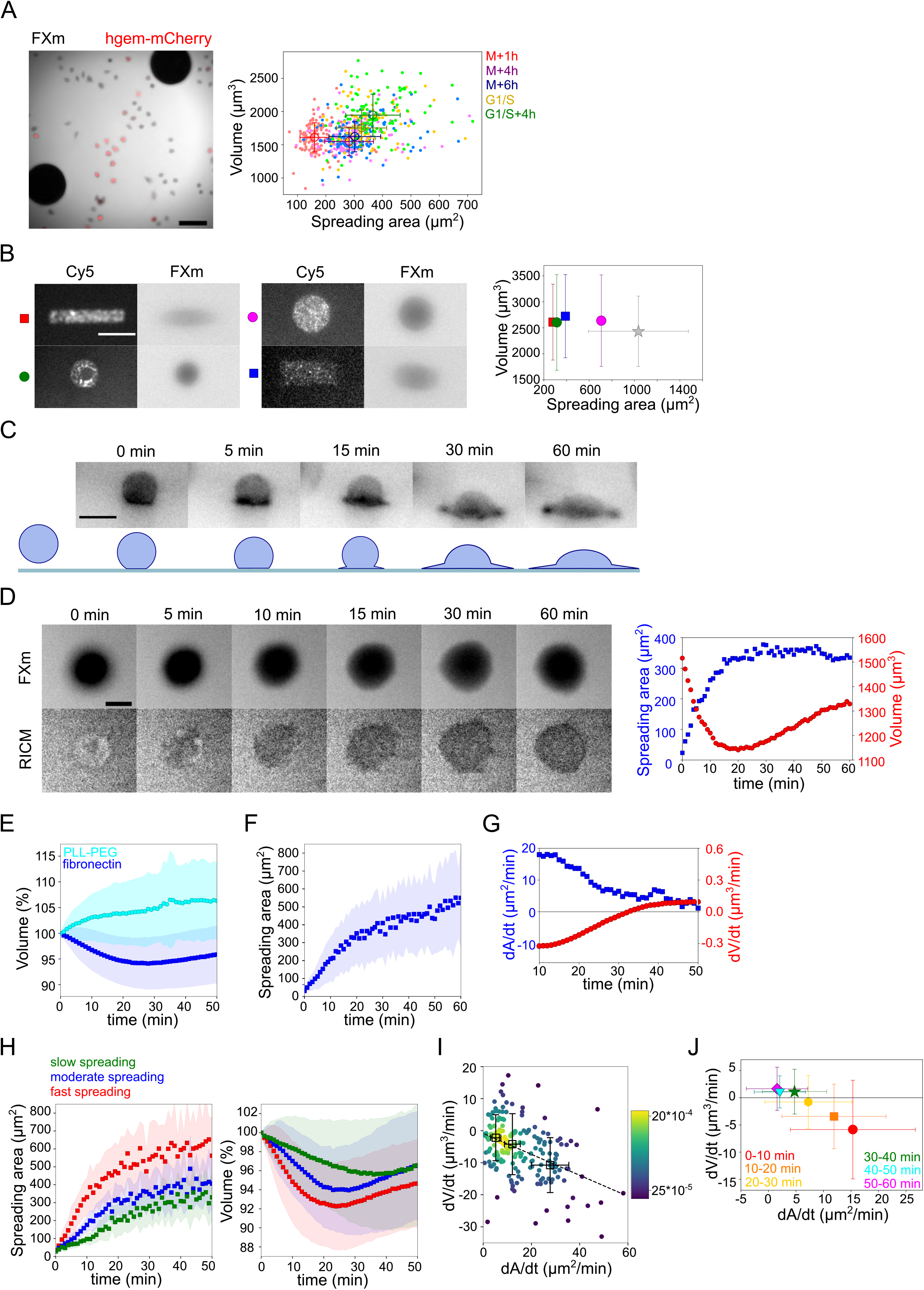
Relation between cell volume and cell spreading. **A. Left:** Composite of FXm in GFP channel and fluorescent image in mCherry channel of HeLa hgem-mCherry cells. Scale bar 100 μm. **Right:** Relation between volume and spreading area of HeLa hgem-mCherry cells at the different cell cycle stages: M+1h (n=131), M+4h (n=131), M+6h (n=131), G1/S (n=99), G1/S+4h (n=92). Error bars represent standard deviation. **B. Left:** Typical images of micropatterns and typical images of cells plated on micropatterns. Scale bar 10 μm. **Right:** Average volume of HeLa Kyoto cells plated on the patterns of different shape and size in comparison with non-patterned cells. Blue: rectangle 30×13 (n=131), red; rectangle 40×7 (n=214), purple: circle, r=15 (n=338), green; circle, r=10 (n=242), grey: non-patterned cells (n=151). Error bars represent standard deviation. **C. Top:** Side view of a HeLa-Lifeact (black) cell spreading on fibronectin-coated glass. Scale bar 20 μm. **Bottom:** Scheme of shape transition during cell spreading. **D. Left:** FXm and RICM imaging of a HeLa Kyoto cell spreading on fibronectin-coated glass. Scale bar 20 μm. **Right:** Volume (red) and spreading area (blue) of cell represented on the left panel. **E.** Average normalized volume of control HeLa Kyoto cells (blue, n=125) spreading on fibronectin-coated glass, or plated on PLL-PEG-coated glass (cyan, n=84). Error bars represent standard deviation. **F.** Average spreading area of control HeLa Kyoto cells (n=125), spreading on fibronectin-coated glass. Error bars represent standard deviation. **G.** Linear derivatives dA/dt (blue) and dV/dt (red) for average spreading area and volume represented on **F** and **E** for sliding window 10 min. **H.** Average normalized volume (**left**) and spreading area (**right**) of control cells divided in 3 categories based on their initial spreading speed: slow (n=42), moderate (n=43), fast (n=42). Error bars represent standard deviation. **I.** Volume flux (dV/dt) of single control HeLa Kyoto cells (n=194) plotted versus their spreading speed (dA/dt) at the first 10 min of spreading. R=-0.41. Error bars represent standard deviation. Color bar indicate kernel density. **J.** Median volume flux (dV/dt) of HeLa Kyoto cells plotted versus median spreading speed (dA/dt) at the different time intervals (n=194). Error bars represent standard deviation.

Previous studies also reported volume loss during cell spreading ^9^. When plated on a fibronectin-coated substrate, HeLa cells showed a transition from a sphere to a half-sphere in about 15 minutes, then continued spreading by extending lamellipodial protrusions (**Figure 1C** and **Movie S1**). We recorded spreading cells, combining FXm to measure volume and Reflection Interference Contrast Microscopy (RICM) to measure spreading area accurately (**Figure 1D** and **Movie S2**). RICM images also showed an initial spreading phase of about 15±10 min until the radius of the contact region equaled that of the cell, corresponding to a hemispheric cap cell shape, which was followed by an extension of lamellipodial protrusions. Cell spreading was accompanied by a small (5% on average) but significant loss of volume, typically occurring during the first 20 min of spreading and followed by a volume increase of about 5%/h, in the range of the expected cell growth for a doubling time of about 20 h **(Figure 1E, F, G)**. The same was observed for cells that had been synchronized by serum starvation, but with a smaller standard deviation (**Figure S1E**). Combining quantitative phase and volume measurement, we found that only cell volume decreased while dry mass remained constant over the few tens of minutes of initial cell spreading (**Figure S1F**), causing a transient density increase (**Figure S1G and 1G**). This suggests a loss of water (and probably small osmolites like ions) from the cell, similar to volume regulatory decrease following a hypo-osmotic shock ^22^. Cells plated on PLL-PEG, instead of fibronectin, did not spread and displayed an increase in the volume of about 7%/h (**Figure 1E**). This result, together with our observation on steady-state spread cells, suggests that the spreading dynamics rather than the final spreading area might be coupled to the loss of volume.

Taking advantage of the intrinsic variability in the cell spreading dynamics, we considered single cell volume and spreading trajectories. We observed that individual spreading cells could display a large range of volume loss (**Figure S1H**). Pooling cells together according to their spreading speed, we observed that faster-spreading cells were losing more volume whereas slow spreading cells lost less volume or did not lose volume at all (**Figure 1H**). To further validate this correlation, we measured the initial spreading speed and plotted it against the rate of volume loss, for individual cells (**Figure 1I**). The graph clearly shows that faster spreading cells also lose volume faster in this initial spreading phase. Spreading speed and volume loss are both slowing down with time (**Figure 1J**), whereas absolute spreading area increases (**Figure S1I**). Overall, these data show that volume loss in spreading cells is a transient phenomenon correlated with the spreading kinetics and not the absolute spreading area.

Early spreading dynamics were shown to strongly depend on the properties of the actomyosin cortex ^23^. Hence, we affected F-actin with a low dose of Latrunculin A (Lat A) which still allowed cell spreading, and myosin with the ROCK inhibitor Y-27632 (Y-27, **Figure 2A**). As expected, we found that Lat A-treated cells spread slower, while Y-27-treated cells spread faster than control cells (**Figure 2B**). Accordingly, Lat A treated cells lost less volume (2-3%) than control cells, while Y-27 treated cells lost more (15%, **Figure 2C, D**). Y-27 treated cells plated on PLL-PEG substrate on which they could not spread, increased their volume like control cells (**Figure S2A**), thereby showing that larger volume loss was not due to the drug treatment itself but was a result of the spreading kinetics in the presence of the drug. This coupling between spreading speed and volume loss was also found to be very similar for other cell types (**Figure S2B-H**), although RPE-1 cells displayed an initial phase of volume increase before eventually losing volume. This initial phase of volume increase was lost upon Y-27 treatment (**Figure S2E, S2H**), suggesting that it was due to induction of contractility through mechanotransduction pathways ^24^. These data together suggest a general effect of spreading kinetics on volume modulation, with a loss of volume reaching up to 20% for fast-spreading cells.

**Figure 2:**
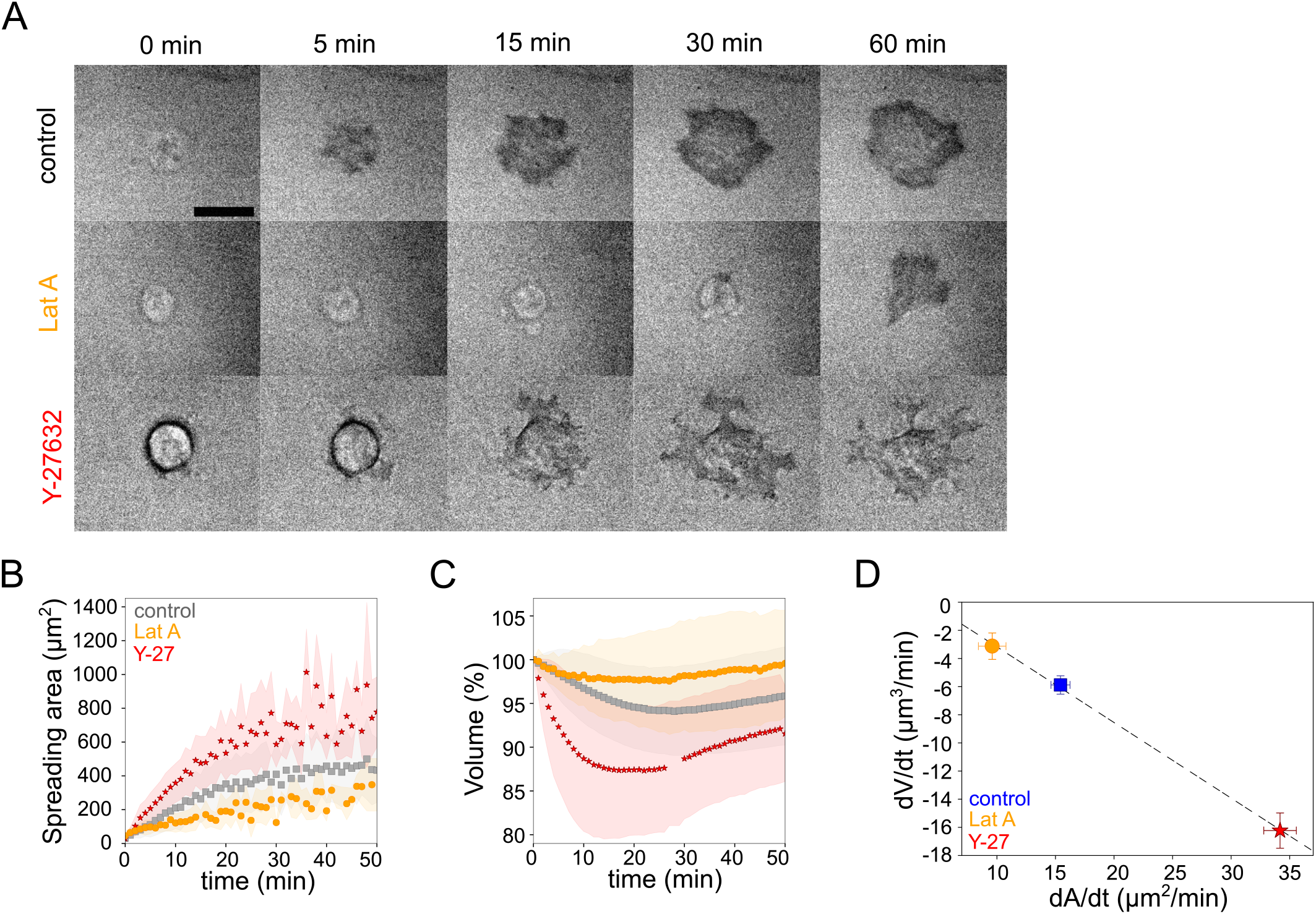
Volume modulation in slow and fast spreading cells. **A.** RICM imaging of control HeLa Kyoto cell or cell treated with 100nM Latrunculin A or with 100 μM Y-27632 spreading on fibronectin-coated glass. Scale bar 20 μm. **B.** Average spreading area of control HeLa Kyoto cells (grey, n=125), cells treated with 100 nM Latrunculin A (orange, n=30) or 100 μM Y-27632 (red, n=98). Error bars represent standard error. **C.** Average normalized volume of control HeLa Kyoto cells (grey, n=125), cells treated with 100 nM Latrunculin A (orange, n=30) or 100 μM Y-27632 (red, n=98). Error bars represent standard error. **D.** Median volume flux (dV/dt) of control (blue, n=194), 100 μM Y-27632 (red, n=121) or 100 nM Latrunculin A (orange, n=30) treated HeLa Kyoto cells plotted versus their spreading speed (dA/dt) at the first 10 min of spreading. Error bars represent standard error.

### The osmotic response of cells follows the Pump and Leak Model (PLM)

Water loss exceeding 1% is considered to be dominated by osmotic volume regulation (see **Supplementary file** and ^21,25^). Volume set point and large volume modulation such as volume regulatory response following osmotic shocks can be accounted for by the general theoretical framework of the ‘pump and leak model’ (PLM, see **Supplementary file** and ^21,26,27^). Briefly, the cell volume is determined by an osmotic balance involving the active pumping of specific ions (sodium and potassium) to compensate for the pressure from impermeant solutes in the cell (**Figure 3A**). The PLM has been verified experimentally on several occasions, mostly with indirect methods for cell volume measurements ^28^. We thus decided to check that we could reproduce these results with our cells. We performed series of osmotic shock experiments while recording cell volume by FXm (**Figure 3B** and **Figure S3A-C** and **Movie S3**). Cells showed the expected response to both hypo and hyperosmotic shocks, with a fast change in volume (less than a minute timescale) followed by a slower adaptation (timescale of minutes) (**Figure 3C** and **Figure S3D**). We also checked, using quantitative phase measure of dry mass, that these fast changes in volume were not accompanied by any change in dry mass and thus corresponded to water (and ion) fluxes, as expected (**Figure S3E**). Because of timescale separation between water flux in the seconds timescales and active ion transport, which takes minutes, upon an osmotic shock, cells first display a passive response corresponding to water fluxes, followed by a slower response due to ion exchanges. The Ponder’s relation ^29^, which relates the relative change in cell volume right after the shock (at timescale of seconds), to the relative difference of osmotic pressure imposed experimentally, corresponds to the passive cell response. The Ponder’s plot showed a very good agreement with previous reports, with a linear relation between the change in volume and the change in osmotic pressure, over a large range of imposed external osmolality (**Figure 3D**). The slope is also similar to previous reports ^30,31^, and corresponds to about 30% of osmotically inactive volume (volume occupied by large molecules or solid components). As shown by others ^30^, we find that the Ponder’s relation does not depend on the integrity of the actin cytoskeleton, as cells treated with Lat A show the same relation (**Figure 3D**). These experiments also allowed us to estimate the bulk modulus of cells defined as 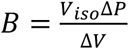, where *V_iso_* is volume in isosmotic state, Δ*V* is volume change, induced by osmotic pressure difference Δ*P* (order of GPa, **Figure S3F**), which is in good agreement with previous measures ^9,32^. These results show both that our cell volume measurements are accurate, even for small volume changes, and that our cells display the expected response to osmotic shocks.

**Figure 3:**
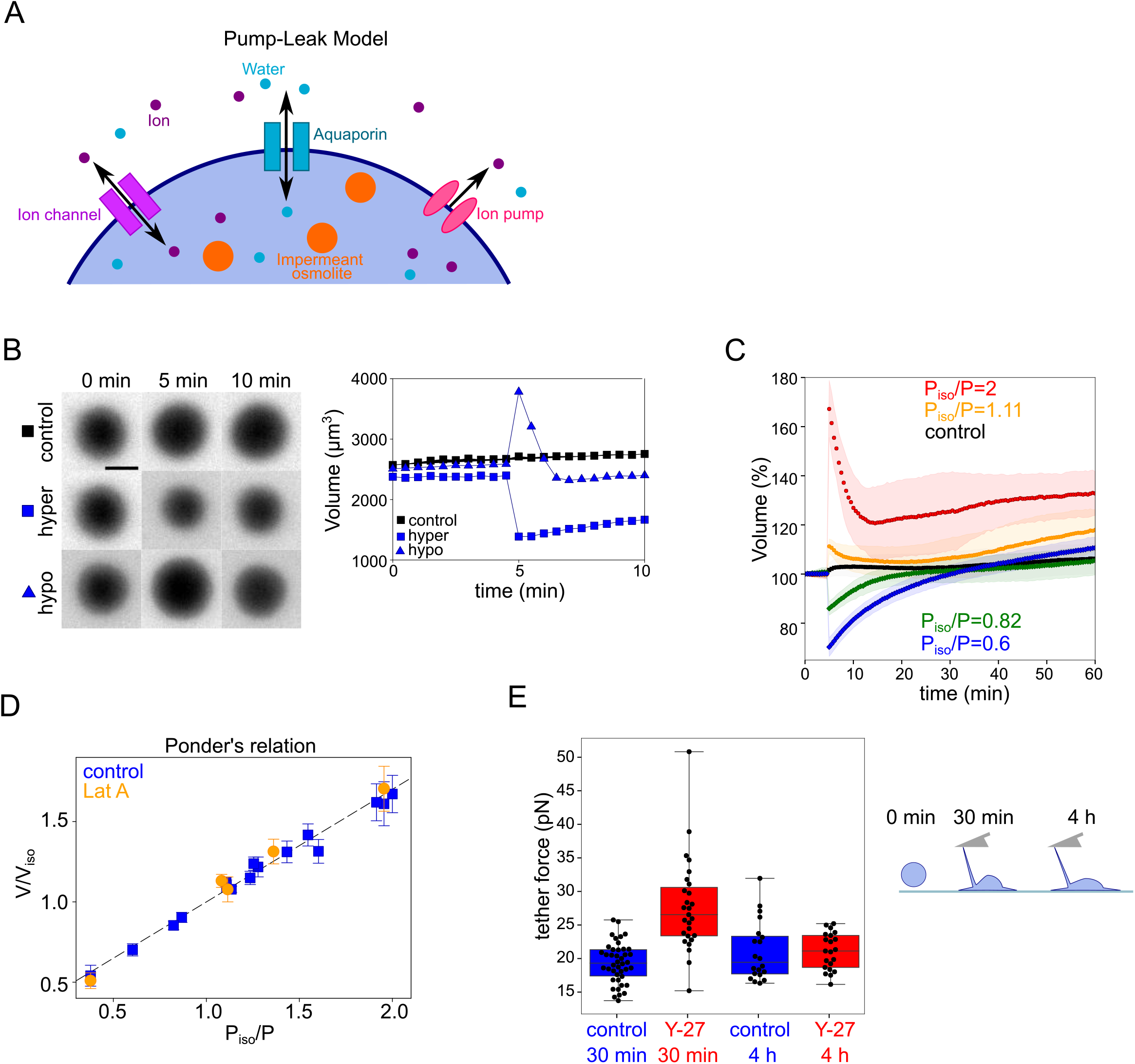
Volume response to osmotic shocks and membrane tension in fast spreading cells. **A**. Schematic of “pump-leak” model. **B. Left:** FXm images of HeLa Kyoto cells exposed to media exchange of same osmolarity, hypertonic and hypotonic. **Right:** Volume of cells represented on the left panel. **C**. Examples of average normalized HeLa Kyoto cells volume response to osmotic shocks of different magnitudes. Average normalized volume of HeLa Kyoto cells in response to osmotic shocks of different magnitudes. Number of cells in the experiments: control P_iso_/P=1 (n=51), P_iso_/P=1.11 (n=30), P_iso_/P=2 (n=17), P_iso_/P=0.82 (n=33), P_iso_/P=0.6 (n=67). **D**. Ponder’s relation for control HeLa Kyoto cells (blue) and HeLa Kyoto cells treated with 2 μM Lat A (orange). Average number of cells in each experiment n~48. Error bars represent standard deviation. **E**. Tether force measurements of control HeLa Kyoto cells (blue) and treated with 100 μM Y-27632 (red) during the first 30-60 min of spreading and 4h after plating.

### Fast spreading cells transiently display an increased apparent membrane tension

Classical PLM does not take into account the cell shape and mechanics. Several additional mechanisms have been proposed to account for the coupling between cell shape and cell volume. A recent model proposed a direct effect of spreading area on cell volume with the assumption that channels and pumps are working differently on the adhered and the free surface of the cell ^12^. Nevertheless, such a model does not predict an effect of the rate of spreading on volume, but rather an effect of the spreading area itself, while our data suggests that the opposite is true in our experiments. The correlation of volume loss with spreading kinetics suggests that the effect on volume could be mediated by the coupling of cell cortex mechanics with the functioning of ion channels and pumps. This has been proposed before, with a simplified version of the PLM, including only one neutral solute transported across the cell with mechanosensitive channels ^10,13,17^. According to this class of models, increased membrane tension would modify the balance of the solutes, which would in turn change cell volume. To test whether fast-spreading cells have a higher membrane tension than slow-spreading cells, we pulled membrane tubes with an Atomic force Spectrometer (AFS) tip following a well-established protocol (see Material and Methods for further details, ^33^). This allows to measure the tether force, which is proportional to an apparent membrane tension. We measured the tether force for cells 30 minutes after plating them on an adhesive substrate and compared the values to steady-state spread cells (4 hours after spreading). We found that cells treated with Y-27, which spread faster than control cells, displayed a higher tether force during the spreading phase, while the force was similar to control cells at steady state (**Figure 3E**), showing that the increase was not due to the drug treatment itself. This experiment suggests an effect of spreading speed on membrane tension. This is consistent with the hypothesis that mechanosensitive processes might modulate cell volume upon fast cell shape changes.

### A mechano-sensitive PLM including a mechano-osmotic coupling predicts the observed relation between spreading speed and volume loss

We note that the one solute model without trapped solute particles proposed before ^17^ does not lead to a result that satisfies both the osmotic balance and the solute transport equation (see details in **Supplementary file**). The osmotic balance in the one solute model implies that the solute concentration should be equal inside and outside the cell, whereas the solute transport fixes the concentration to a value set by the pumps and channels. Thus, the osmotic balance and solute transport equations do not lead to a consistent solution. We thus engaged in proposing a modified model (see the full model in **Supplementary file**), to combine PLM with cell mechanics and shape, assuming that ion channels and pumps can be affected by membrane tension, as demonstrated multiple times by others ^34^. The model also includes cell growth, with the experimentally measured rate, playing a significant role in long timescales.

In brief, in this new model (see details in **Supplementary file**), the tension dependence of volume is through the mechano-sensitivity of the ion channels and pumps. In the linear regime, we assume that small changes in tension lead to a small change in ion transport rates so that the volume change is proportional to the change in tension. The equation for change in volume reads

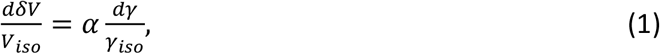

where *δV* = *V*(*t*) – *V_iso_*(1 + *gt*), with *g* and *t* being the growth rate of the cell and time respectively. If the mechano-sensitivity parameter *α* is negative then the volume will decrease upon an increase in the tension. We explicitly evaluate the mechano-sensitivity parameter *α*, by analyzing a model of ion transport with three ion species - sodium, potassium, and chloride. We find that the sign and magnitude of *α* depend on the mechano-sensitivity of the potassium and sodium channels, and on the ion concentrations before spreading. For physiological values of parameters found in the literature (see values in **Supplementary file**), we expect *α* to be negative. To relate tension variations to the rate of cell spreading, we model surface tension using a Maxwell fluid model, with a relaxation timescale *τ* and elastic modulus *k*, driven by the rate change of total surface area. The tension dynamics reads

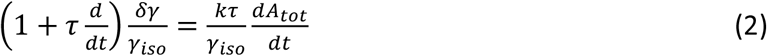

The elastic modulus characterizes the short time elastic response while the relaxation timescale accounts for the existence of tension homeostasis mechanisms that have a longer response time. During cell spreading, the total surface area will increase leading to a spreading rate-dependent increase in tension, which will relax back to the homeostatic value, in agreement with the tether pulling experiments reported in **Figure 3E**. To estimate the total surface area, we take the cell shape to be that of a spherical cap (**Movie S4;** we discuss in more details the possible shape approximations and their relation to the measured spreading area in the **Supplementary file**). Combining the tension dynamics in **Equation (2)** with tension/volume coupling in **Equation (1)** leads to the following effective viscoelastic model for volume dynamics driven by a change in the total area,

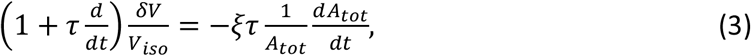

where the effective elastic modulus *ξ* = –(*A_tot_kα*)/*γ_iso_* is proportional to the effective elasticity of the membrane *k* and to the magnitude of the mechano-sensitivity parameter *α* relating volume loss to the tension increase. *ξ* is also inversely proportional to the surface tension. The total area itself depends on the volume, we can write **Equation (3)** as

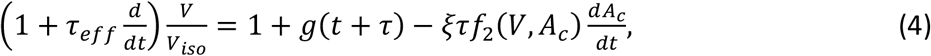

where *f*_1_(*V, A_c_*) and *f*_2_(*V, A_c_*) are functions that are given by the geometry of the cell, which relate the change in total area to the change in volume and change in contact area respectively. The effective relaxation timescale of the volume and tension *τ_eff_* = *τ*(1 + *ξf*_1_(*V, A_c_*)*V_iso_*) is proportional to *τ* but also involves *ξ* and a function that depends on cell shape and size. This new model has thus two main fitting parameters *ξ* and *τ* that relate the spreading area to the volume change. These parameters allowed us to fit the various experimental data and their values are discussed in more details below. Importantly, this simple extension of the PLM predicts the observed proportionality between volume loss and the speed of spreading, and no dependency on the absolute cell spreading area (**Equation (3)**, **Figure 1I** and **2C, D**). We conclude that this new model thus constitutes a robust implementation of a membrane tension homeostasis mechanism within the PLM framework and we propose to call it the mechano-sensitive PLM.

### Fitting the spreading and volume data with the mechano-sensitive PLM

To verify the main assumptions of the model, such as the timescales of water and ion fluxes, we performed a detailed characterization of the cell response to osmotic shocks. We first made high time resolution recordings of cell swelling and shrinking upon a change in the external osmolarity (**Figure 4A** and **S4A, B** and **Movie S5**). The change of volume occurred in a timescale of seconds, as expected. These experiments provided the rate of cell water entry and exit as a function of the difference in osmotic pressure (**Figure 4B**). This allowed us to estimate the hydraulic conductivity (**Supplementary file**, **Figure S4C**), which appeared smaller for hyper-osmotic shocks than for hypo-osmotic shocks, as reported previously ^35,36^, although the reason for this difference is not understood. We next characterized the longer, minutes timescale of volume adaptation (**Figure 4C**). It showed that volume adapted faster for larger shocks. At the level of individual cells, the response was quite homogenous for the recovery from hypertonic shocks, while there was a higher cell-cell variability during recovery from hypotonic shocks (**Figure S4D**), with cells showing only partial recovery, especially for large hypo-osmotic shocks. Despite these complex single-cell behaviors, these experiments provide clear evidence, as well known from decades of studies of this phenomenon, of a volume regulation mechanism on the timescale of minutes, setting the typical timescale for ion fluxes. These two timescales, seconds for water flows through the membrane and minutes for ion fluxes, are basic assumptions of the PLM model verified by our experiments. Importantly, the rate of volume change observed for small shocks is similar to the rate of volume change during cell spreading experiments (about 10 μm^3^/min). This justifies the use of a mechano-sensitive PLM to explain the cell spreading data. We thus performed a fit of our experimental data using the mechano-sensitive PLM model. We used the three groups of control cells defined in **Figure 1H**, sorted based on spreading speed during the first 10 minutes. The spreading parameters were extracted from the experimental spreading data, and the model allowed a satisfactory fit of the experimental volume data (**Figure 4D**).

**Figure 4:**
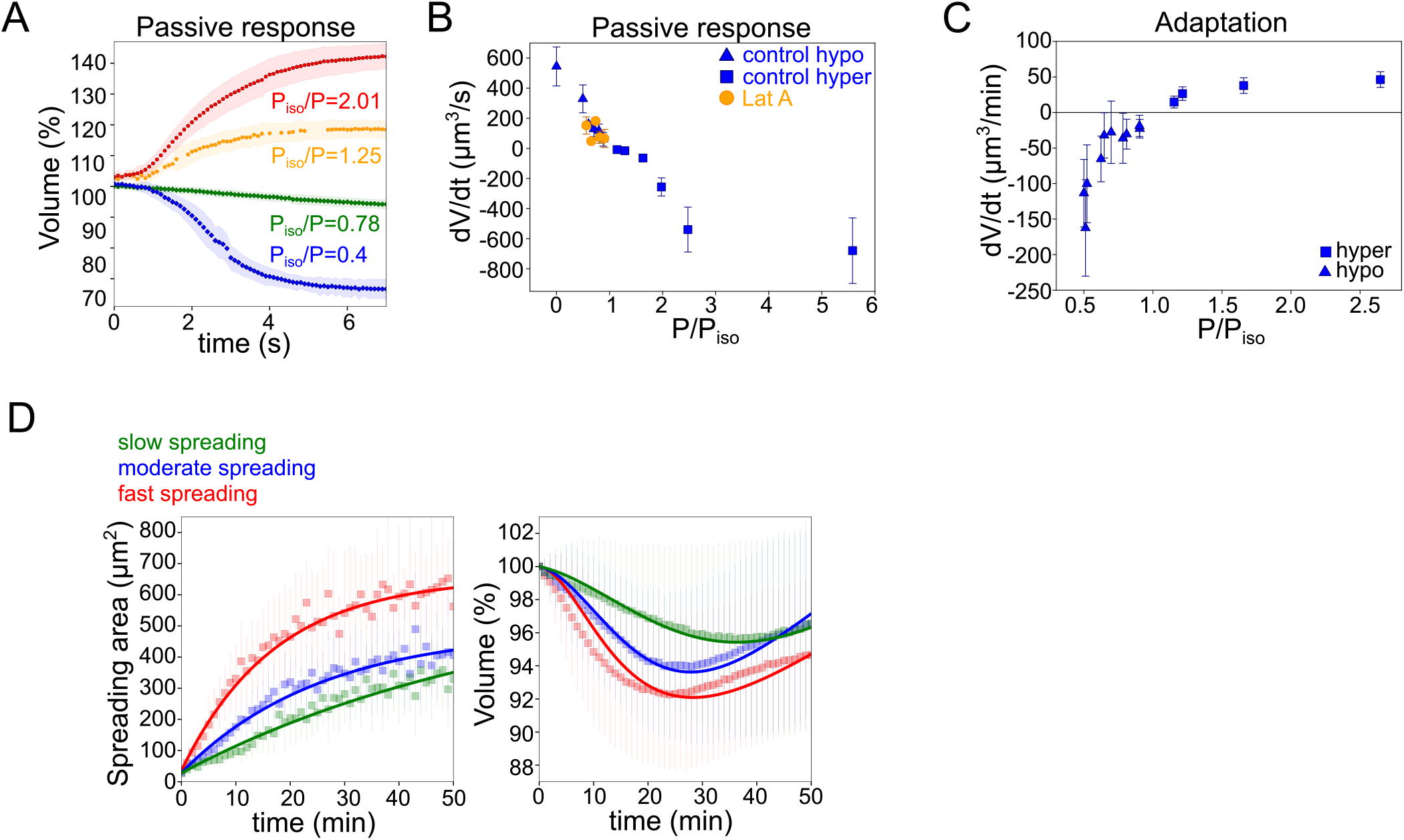
Timescales of the osmotic response and fitting of the cell spreading data with the mechanosensitive PLM. **A.** Average normalized volume of HeLa Kyoto cells during initial response to osmotic shocks of different magnitudes measured with high time resolution. Number of cells in the experiments: control P_iso_/P=1.25 (n=13), P_iso_/P=2.01 (n=10), P_iso_/P=0.78 (n=15), P_iso_/P=0.4 (n=17). Error bars represent standard deviation. **B.** Average volume flux in HeLa Kyoto cells during initial response to osmotic shocks of different magnitudes. Average number of cells in each experiment n~12. Error bars represent standard deviation. **C.** Average volume flux in HeLa Kyoto cells during regulatory volume adaptation. Average number of cells in each experiment n~48. Error bars represent standard deviation. **D.** Fits from the model using best fit parameters on average normalized volume (**left**) and spreading area (**right**) of control cells divided in 3 categories represented in Figure **1H**.

Because the mechano-sensitive PLM assumes a coupling between membrane tension and ion fluxes, how much volume is lost by a cell during spreading depends on whether the cell deforms in a rather elastic or viscous regime. The transition between these regimes is defined by the relative values of the spreading rate *τ*_a_ and the effective tension relaxation time - if the spreading rate is faster than the relaxation time, the cell deforms in a rather elastic regime, and as a result, the membrane gets tensed and the cell loses volume. The effective relaxation time depends on the two main fitting parameters, the bare tension relaxation timescale *τ* (which varies in the minutes to tens of minutes timescale) and the stiffness *ξ* (which varies around one). When fitting the three classes of fast, intermediate and slow-spreading cells, we found that the values of the fitting parameters (**Table III** in **Supplementary file**), do vary significantly for the three classes. However, this variation could not explain the difference in volume loss (see **Supplementary file**) which must therefore be attributed to the difference in spreading speed. We conclude that our mechano-sensitive PLM not only captures properly the coupling of spreading kinetics on volume modulation but that the parameter fitting suggests that the key ingredient of the model, the finite response time of the mechano-osmotic feedback, might be the cause of the volume loss in fast-spreading cells.

### Increased membrane tension and volume loss during fast deformation is due to the actin cortex

We then asked what could be the origin of the increase in surface tension during fast cell spreading. We first evaluated the total amount of cell membrane available. We exposed cells to distilled water and first imaged actin and membrane staining. It showed a rapid full unfolding of membrane reservoirs (**Figure 5A**) before the cell exploded. We then used propidium iodide to identify the timing of plasma membrane rupture (**Figure 5B**). We found that on average, the plasma membrane ruptured when cells reached 5.7 times their initial volume, which corresponds to an excess of membrane surface area of about 3.3 times, in accordance with previous measures ^37,38^.

**Figure 5:**
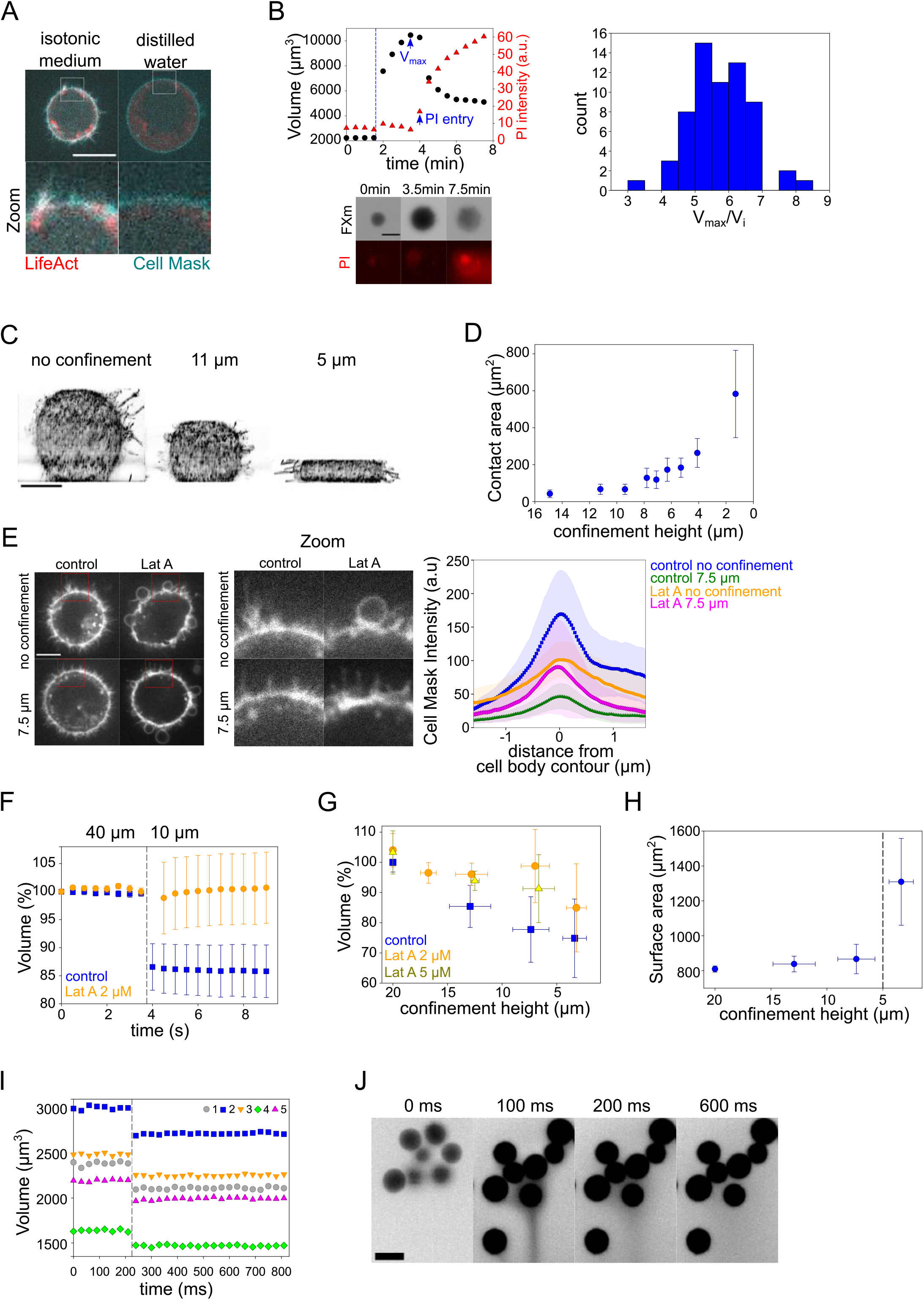
Measure of total plasma membrane surface area and response of cells to fast compression. **A.** Z-plane of HeLa LifeAct-mcherry (red) cell before and after addition of distilled water, cell membrane is stained with CellMask Green (cyan). Scale bar 10 μm. **B. Left top:** Volume (black) and propidium iodide (PI) intensity of single HeLa Kyoto cell exposed to distilled water. Dashed line indicates the time of distilled water addition. Reaching of maximum cell volume is followed by cell membrane rupture, volume decrease and PI entry into the cell. **Left bottom:** Corresponding FXm images and propidium iodide (PI) staining. **Right:** Distribution of ratio between maximum volume cells reach before bursting induced by exposure of distilled water and their initial volume. **C**. 3D-membrane reconstruction of HeLa expressing MyrPalm-GFP (black) cells cell shape under different confinement heights, side view. Scale bar 10 μm. **D**. Contact area with bottom glass substrate of Hela Kyoto cells under different confinement heights. Average number of cells in each experiment n~79. Error bars represent standard deviation. **E. Left:** Z-plane of control and 2 μM Lat A treated HeLa-MYH9-GFP-LifeAct-mcherry, cells under 20 μm and 7.6 μm confinement heights. Cell membrane is stained with CellMask Far Red (white). Scale bar 10 μm. **Right:** Average CellMask intensity plotted versus distance from cell body contour on the middle Z-plane of HeLa-MYH9-GFP-LifeAct-mcherry cells. Number of cells in each condition n=10. Error bars represent standard deviation. **F.** Average normalized volume of control (blue, n=48) and 2 μM Lat A treated (orange, n=32) HeLa Kyoto cells during dynamic confinement experiment. Dashed line indicates the moment of confinement. Error bars represent standard deviation. **G.** Average normalized volume of HeLa Kyoto cells (blue) and cells treated with Lat A 2 μM (orange) or 5 μM (yellow) under different confinement heights. Each data point represents an average of N~9 experiments; each experiment contains n~137 individual cells. Error bars represent standard deviation. **H**. Projected surface (computed from volume represented on panel **Figure 5G**) of HeLa Kyoto cells under different confinement heights. Dashed line indicates the confinement height that corresponds to blebs appearance. Error bars represent standard deviation. **I.** Volume of single HeLa Kyoto cells during dynamic confinement experiment. Dashed line indicates the moment of confinement. **J**. FXm images of HeLa Kyoto cells during dynamic confinement experiment taken with high NA objective.

It means that cell have a very large excess of membrane surface area and that membrane tension could not arise from a limitation in membrane availability. Nevertheless, the plasma membrane being bound to the underlying cytoskeleton, its restricted unfolding could generate an increase in tension depending on the rate of cell deformation. We thus investigated fast cell deformations.

To impose fast (less than a second timescale) deformation on the cell, we used our previously developed cell confiner ^39^. This device can impose a precise height on cells and thus gives access to a large range of deformations (**Figure 5C** and **S5A** and **Movie S6**). RICM measure of the cell contact area showed a range of spreading similar to what was observed during spontaneous cell spreading (**Figure 5D** and **S5B**). In addition, imaging of the plasma membrane showed that confinement below 10 μm induced a clear loss of membrane folds and reservoirs (**Figure 5E** images and **Movie S7**), while treatment with Lat A induced the formation of large membrane blebs and less extension of the cell diameter upon confinement (**Figure 5E** graph and **S5C** and **Movie S7**). This suggests that cell confinement, like hypo-osmotic shocks, induces membrane reservoir unfolding, and that Lat A treatment, by reducing the membrane anchorage and causing bleb formation, reduces the surface expansion following confinement. FXm volume measurement combined with confinement showed a strong loss of volume of confined control cells, while Lat A treated cells kept a constant volume (**Figure 5F** and **S5D** for a control showing for both treatments the decrease in FXm background intensity corresponding to the confiner height; and **Movie S8**). In control cells, stronger confinement led to larger volume loss, while Lat A treated cells showed no significant volume loss except for the lowest confinement height (**Figure 5G**). The loss of volume in control cells corresponded to a deformation at an almost constant surface area (**Figure 5H**, calculated from the volume, see **Supplementary file**). Below 5 μm height, the cell surface significantly increased, which also corresponded to the formation of large blebs (**Figure S5E**). This loss of volume induced by fast confinement was also found in other cell types (**Figure S5F, G**) and was also previously observed in confined *Dictyostelium* cells ^40^. Overall, these experiments show that fast imposed cell deformation induces an actin-dependent loss of volume (up to 30%), at almost constant surface area.

To better estimate the speed of deformation imposed by the confiner, we imaged at high frame rate during the confinement process. It showed that, even with a time-lapse of 30 ms, the volume loss happened between two consecutive frames (**Figure 5I** and **S5H, I** and **Movie S9**). Only volume is lost and not dry mass (**Figure S5J**), which suggests that only water and probably small solutes are lost. Nevertheless, the speed of volume change is not compatible with our mechano-sensitive PLM, as in this model, volume loss occurs in the minutes timescale due to changes in ion transport rates. Fast imaging of the fluorescent medium surrounding the cells used for FXm indeed showed a transient appearance of streams of darker fluid (non labelled, thus coming from the cells) emanating from confined groups of cells (**Figure 5J** and **Movie S9**).), likely corresponding to the expelled water and osmolites. Overall, these confinement experiments suggest that, although at this timescale of milliseconds, the mechanism of volume loss very likely differs from the context of spontaneous cell spreading, it is also induced by an increase in membrane tension, and requires the presence of the actin cytoskeleton.

### Volume loss upon fast cell deformation depends on branched actin and on changes in ion fluxes

Because branched actin was shown to more specifically interact with the plasma membrane ^33,41^ and modulate membrane tension, we used cells treated with the Arp2/3 complex inhibitor CK-666, and combined the treatment with Y-27 to induce fast spreading. We found that CK-666 treatment alone induced both a slower spreading and lower volume loss (2-3%, **Figure 6A, B**), similar to the low Lat A treatment (**Figure 2B, C**), which was well fitted by the mechanosensitive PLM (fits on **Figure 6A, B**). Treatment with Y-27 increased the spreading speed of CK-666 treated cells, but the volume did not decrease in this fast-spreading condition (**Figure 6C, D**). The mechano-sensitive PLM could fit these data by adjusting the parameter coupling surface tension to the change of activity of ion pumps (**Table IV** in **Supplementary file**). To directly test the role of ion fluxes in the volume loss, we targeted two main players: first, stretch-activated calcium channels (including Piezo), using gadolinium chloride (GdCl_3_) and second NHE1, the sodium/proton exchanger, using EIPA. Treatment with GdCl_3_ led to an increase in volume loss (from 5 to 8%), which could be fully accounted for by the increase in cell spreading speed (**Figure 6E, F**) and well fitted by the mechano-sensitive PLM. This effect can be explained by the known effect of calcium on actomyosin motors activation ^42^. In the context of cell spreading, the opening of these channels would let calcium in the cell, leading to activation of actomyosin contractility, which is known to act against spreading ^43^. The absence of disruption of the coupling between spreading speed and volume loss is also expected since calcium is a potent second messenger, but its concentration in the cell is not compatible with direct volume regulation. On the other hand, cells treated with EIPA, while spreading slightly faster than control cells, lost less volume (**Figure 6G, H**). Combining EIPA with Y-27 showed that, despite a spreading speed comparable to Y-27 treated cells, NHE1 inhibition fully prevented volume loss (**Figure 6G, H**). Inhibition of NHE1, which is known to affect ion transport, is thus also preventing volume loss during fast spreading. This is consistent with the role of a change in ion fluxes in the mechano-sensitive PLM.

**Figure 6:**
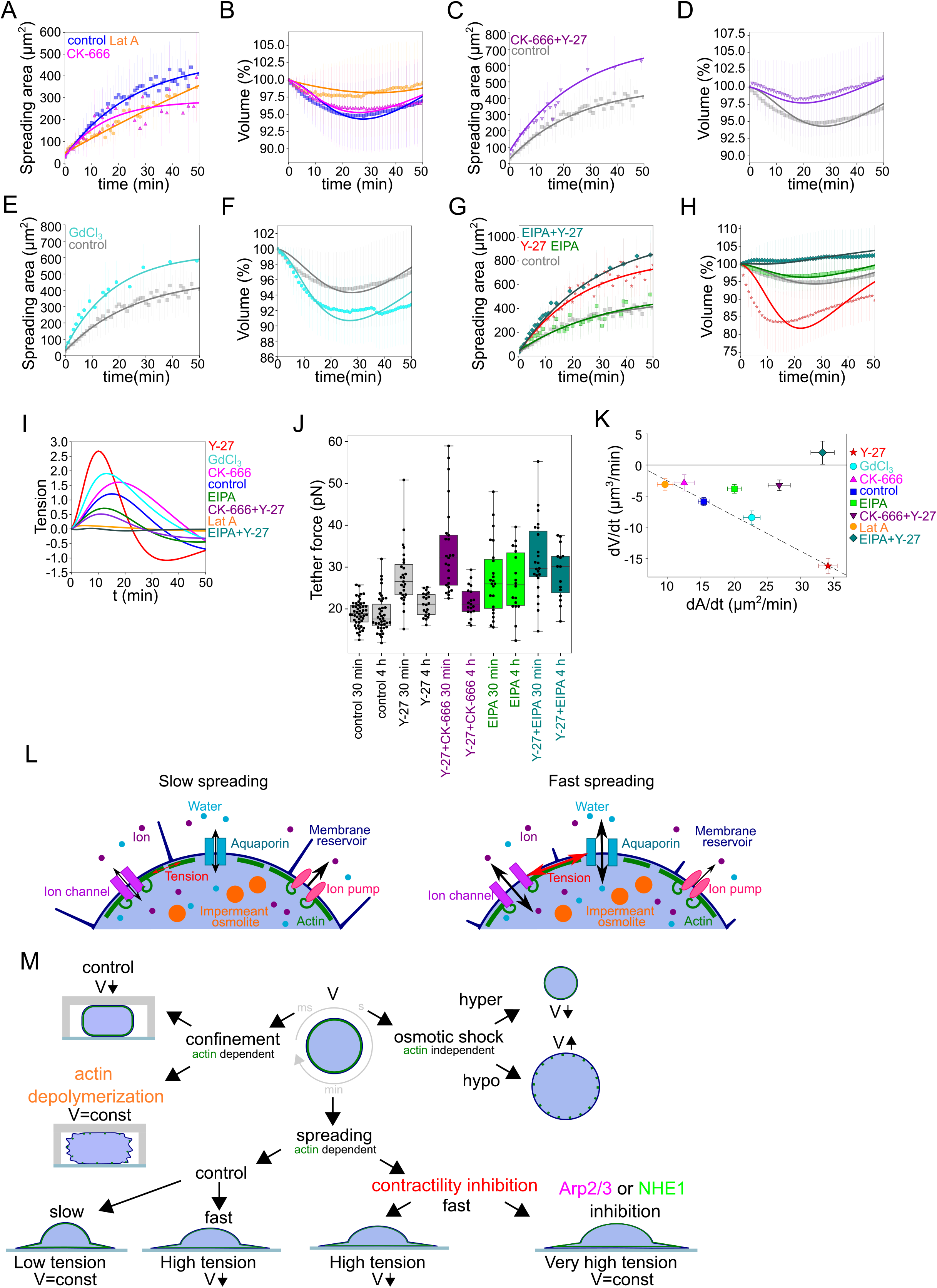
Perturbations of branched actin networks and ion fluxes and associated changes in volume and surface tension of spreading cells. **A.** Two parameter fits for the spreading kinetics using the exponential saturation anzatz (see text) on average area of control cells (blue, n=73), 100 nM Latrunculin A (orange, n=30) or 100 μM CK-666 (magenta, n=37) treated. Error bars represent standard deviation. **B.** Fits from the model using best fit parameters on average normalized volume of control cells (blue, n=73), 100 nM Latrunculin A (orange, n=30) or 100 μM CK-666 (magenta, n=37) treated. Error bars represent standard deviation. **C.** Two parameter fits for the spreading kinetics using the exponential saturation anzatz (see text) on average area of control cells (grey, n=73) or combination of 100 μM CK-666 and 100 μM Y-27632 (violet, n=24) treated. Error bars represent standard deviation. **D.** Fits from the model using best fit parameters on average normalized volume of control cells (grey, n=73) or combination of 100 μM CK-666 and 100 μM Y-27632 (violet, n=24) treated. Error bars represent standard deviation. **E.** Two parameter fits for the spreading kinetics using the exponential saturation anzatz (see text) on average area of control cells (grey, n=73) or combination of 100 μM GdCl_3_ (cyan, n=30) treated. Error bars represent standard deviation. **F.** Fits from the model using best fit parameters on average normalized volume of control cells (grey, n=73) or combination of 100 μM GdCl_3_ (cyan, n=30) treated. Error bars represent standard deviation. **G.** Two parameter fits for the spreading kinetics using the exponential saturation anzatz (see text) on average area of control cells (grey, n=73), 100 μM Y-27632 (red, n=21), 50 μM EIPA (green, n=73), or combination of 50 μM EIPA and 100 μM Y-27632 (dark cyan, n=30) treated. Error bars represent standard deviation. **H.** Fits from the model using best fit parameters on average normalized volume of control cells (grey, n=73), 100 μM Y-27632 (red, n=21), 50 μM EIPA (green, n=73), or combination of 50 μM EIPA and 100 μM Y-27632 (dark cyan, n=30) treated. Error bars represent standard deviation. **I.** Predicted by model, plots for difference between tension without mechano-osmotic coupling (for *ξ*=0 and 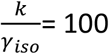) and tension with mechano-osmotic coupling (for fitted *ξ* and 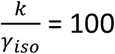) **J**. Tether force measurements of control HeLa Kyoto cells (grey), treated with Y-27632 (grey), CK-666+Y-27 (purple), EIPA (green), CK-666+Y-27 (dark cyan) during the first 30-60 min of spreading and 4h after plating. **K.** Volume flux (dV/dt) of single control HeLa Kyoto cells (n=194), treated with Lat A (n=41), CK-666 (n=54), Y-27 (n=121), EIPA (n=117), GdCl_3_ (n=53), CK-666+Y-27 (n=74), EIPA+Y-27 (n=50) plotted versus their spreading speed (dA/dt) at the first 10 min of spreading. Error bars represent standard error. **L.** Scheme of mechano-sensistive “pump-leak” model. **M.** Scheme representing cell volume regulation in response to deformations

### The mechano-osmotic coupling moderates the membrane tension increase in fast spreading cells, acting as a membrane tension homeostasis mechanism

The mechano-sensitive PLM predicts that inhibition of volume loss in fast-spreading cells should affect membrane tension during the spreading phase (**Figure 6I**). This prediction was tested by tether pulling experiments (**Figure 6J**). These experiments showed that, upon double treatment with CK-666, apparent membrane tension increased transiently even higher than with the Y-27 treatment alone during spreading, while it had similar values in steady-state spread cells (4 h after spreading). Similarly, combined EIPA and Y-27 treated cells showed higher tension than Y-27 or EIPA alone. Tension was highest during early spreading compared to steady-state spread cells, suggesting that the increase was due to spreading and not to the drug treatments alone, even though EIPA alone also add an effect on steady-state tension. This shows that, in these cells, the coupling between membrane tension and volume regulation is lost, and that fast spreading in the absence of volume loss induces higher tension increase (**Figure 6J, K**). Together these experiments confirm the validity of our mechanosensitive PLM. They also support the existence of a membrane tension homeostasis mechanism that reduces the extent of changes in membrane tension upon fast cell shape changes by modulating the relative contribution of surface expansion and volume loss.

## Discussion

### A mechano-osmotic coupling leads to volume loss in fast spreading cells

Our detailed characterization of cell volume during cell deformation upon cell spreading revealed that, while cell volume is not related to the steady-state shape of the cell, it is modulated by the rate of cell shape change. We propose that this is due to a coupling between cell membrane tension and rates of ion fluxes (**Figure 6L**). An extension of the classical pump and leak model (PLM) including this coupling can account for our observations of cell volume during cell spreading. Measures of membrane tension during spreading and at steady state under a variety of conditions confirmed that fast spreading is associated with a transient increase in membrane tension and that preventing volume modulation leads to even higher membrane tension, as predicted by the model. Together, these experiments and this model suggest the existence of a mechano-osmotic coupling at the level of the cell membrane, which acts as a membrane tension homeostasis mechanism by reducing membrane tension changes upon fast cell deformation (**Figure 6M**).

### Volume loss in spreading cells is not correlated with traction forces

As cells spread on a substrate, they also exert forces on it, called traction forces. Because traction forces acting on focal adhesions are well-known mechano-transducer elements, leading to a large number of signaling cascades, they could also contribute to change ion fluxes and modulate the cell volume. We thus performed traction force measurements during cell spreading experiments (**Figure S6A**). These measures showed that, as expected, cells treated with Y-27, which spread faster and lose more volume than control cells, displayed much lower traction forces during spreading. Cells treated with CK-666, which spread more slowly and lose less volume, also displayed smaller traction forces during spreading, and cells treated with both CK-666 and Y-27, which spread fast without volume loss also show low traction forces. Overall, these experiments suggest that traction forces are not related to the amount of volume loss.

### The role of membrane binding to the actin cortex in inducing membrane tension and volume loss in fast spreading cells

A central hypothesis in the model is that the physical coupling between the actin cortex and the cell membrane leads to an increase in membrane tension when the rate of deformation is faster than the relaxation time of the actin cortex and membrane ensemble. To verify this hypothesis, we performed membrane tether experiments in various conditions, during spreading and at steady state (**Figure 6J**). We also imaged membrane structures in the 2D confinement experiments (**Figure 5E**), which were well suited for such imaging, showing that the plasma membrane was generally extended due to the confinement, but that this was not happening when the actin cortex was perturbed upon Lat A treatment. Membrane-to-cortex attachment is at least partly mediated by proteins of the ERM family ^14^. Thus, we performed an additional experiment using an Ezrin inhibitor (NSC) and monitored cell volume during spreading. We found that while spreading was similar or even slightly faster during the initial phase, treated cells lost less volume than control cells **(Fig. S6B-D)**, consistent with a role of cortex/membrane coupling in mediating the effect of spreading kinetics on volume loss. To further investigate the ultrastructure of the cell cortex during spreading, we unroofed Hela cells after 30 minutes spreading on a fibronectin-coated substrate. These experiments confirmed the different extent of spreading in the various conditions assayed, and the perturbation of branched actin in the CK-666 treated cells (**Figure S6E**). Membrane folds and structures such as clathrin-coated pits and caveolae were present in all conditions. Although their number and degree of curvature did not change significantly, the different populations of caveolae appeared hard to quantify and compare on spreading cells without underestimating the amount of flat caveolae. We conclude that, while our biophysical measures and fluorescence imaging gave a clear indication of changes in membrane tension and folding state during large cell deformations, further investigations are needed to precisely describe the change in the state of the membrane and its association to the actin cortex in this context.

### The sign of the volume change upon fast cell deformation

Although the effect of mechanosensitive ion channels on volume has been discussed before for simplified systems considering one or two solutes ^9,10,12,17^, the relation between mechanosensitivity of ion channels and pumps and volume change is far from obvious. The sign of the volume change upon an increase in tension depends on whether the contribution of ions to the osmotic pressure increases or decreases. Since the cell is always osmotically balanced, if the concentration of the ions inside the cell decreases, the concentration of the trapped molecules should increase by decreasing the volume such that the cell osmotic pressure stays constant and vice-versa. For instance, an increase in sodium channel conductance upon an increase in tension leads to an increase in volume, whereas an increase in potassium conductance upon an increase in tension leads to a decrease in volume. Using a detailed model of ion transport, we show that for physiological values of parameters, as observed in experiments, the volume is indeed expected to decrease upon an increase in tension.

### Values of fitting parameters for the mechanosensitive PLM suggest a role for branched actin in modulating ion fluxes

This new mechanosensitive PLM gives a Maxwell viscoelastic model for the volume with two fitting parameters, *ξ* the effective stiffness, and the bare relaxation timescale *τ*_eff_. In most cases, we get a good quantitative fit for cells treated with different drugs that perturb the cytoskeleton and the ion channels (**Supplementary file**). For the Y-27 and Lat A treated cells the fits only qualitatively capture the temporal dynamics of the volume. One of the reasons for an imperfect fit for these two drugs could be the failure of the spherical cap approximation used to estimate the surface area (see more discussion on the shape estimates and the parameters used for cell surface area in the model, **Supplementary file**). Control cells, and cells treated with Y-27, EIPA, CK-666 show less than 30% variation in the value of *ξ*, implying that most of the volume loss is explained by differences in spreading speed. Cells treated with GdCl_3_ show a larger decrease of 60 % but stay in the same range of parameters (and they are close to the same line in the dV/dt versus dA/dt summary graph shown in **Figure 6K**). However, for the cells treated with Y-27+EIPA and Y-27+CK-666, *ξ* decreases by an order of magnitude, leading to low volume loss even though the cells are spreading fast (**Table IV** in **Supplementary file**). This decrease of *ξ* could be either due to a decrease in the elasticity of the membrane or due to the decrease in the value of the mechano-sensitivity parameter. Spreading experiments show that, for both Y-27+CK-666 and Y-27+EIPA, membrane tension reaches the highest values. This means that, in both cases, spreading is still inducing an increase in membrane tension, and the absence of volume loss reinforces the effect on membrane tension. It suggests that the elasticity parameter is not affected but rather the volume-tension electromechanical coupling. This could mean that, unexpectedly, branched actin networks are specifically required for this coupling. This could be due to a direct association of branched actin with ion channels and pumps ^44,45^.

### Volume loss in ultra-fast deforming cells

Within the PLM framework, for a given osmolarity of an external medium, the cell volume may change either due to a change in hydrostatic or osmotic pressure. Fast compression can increase the cortical tension, which can cause an increase in hydrostatic pressure of the cell.

However, the maximum hydrostatic pressure in the cell before the membrane detaches from the cortex is of the order of 10^2^ Pa, thus producing no direct effect on the cell volume, as discussed before ^21,25^. Hence, the observed volume loss of ten to fifteen percent can only be due to a change in the osmolarity of the cell, and not to a change in hydrostatic pressure. For ion transport to take place at timescales of milliseconds, the transport rates of channels and pumps would need to increase by four orders of magnitude. Such an increase can be easily attained if the high tension upon compression leads to transient formation of pores in the plasma membrane (observed in spreading GUVs ^46^). If these pores are small enough to allow for free ion transport but do not let the larger molecules trapped in the cell pass through (which should be the case since the dry mass was found to remain constant), the cell volume will increase rather than decrease (a consequence of the Donnan effect ^47^). The formation of pores thus cannot explain our observations. Another mechanism that may lead to volume decrease upon compression without losing the trapped osmolites requires a selective increase of the ion conductance upon compression, but by orders of magnitude. Whether the ion conductance can increase by four orders of magnitude by mechanical stretching requires further investigation. Finally, it is also possible that due to its poroelastic nature ^8,48^ the cytoplasm behaves as a gel-like structure, and that water and osmolites are pressed out of the cell upon confinement, without changing the osmotic balance nor the dry mass ^49^. In conclusion, confinement experiments confirm that fast deformation is associated to volume loss in an actin-dependent way, also suggesting a coupling between cell mechanics and volume regulation. However, they are hard to fully interpret in physical terms. This means that such a simple experiment as squeezing a cell cannot yet be understood with the current general knowledge on cell biophysics, pointing to a need for further investigations of the physics of large cell deformations. Such deformations are likely to occur in physiological contexts such as circulation of white blood cells and circulating cancer cells through small capillaries and may lead to volume change as was shown *in vitro* ^7^.

### Volume fluctuations in fast migrating immune cells can be explained by the mechanosensitive PLM

While our mechano-sensitive PLM might be limited in the interpretation of cell deformations occurring below the second timescale, it captures well the larger timescales, based on a modulation of ion fluxes by membrane tension. Such timescales correspond to deformations that cells experience for example as they migrate through dense tissues. This implies that migrating cells might display volume fluctuations. To test this prediction, we used a classical cell migration assay with fast-moving bone-marrow-derived dendritic cells from mice embedded in a collagen gel ^50^. The collagen gel mixed with fluorescent dextran was assembled inside a cell volume measurement chamber (**Figure S6F** and **Movie S10**). Because of the low fraction of collagen in the solution and the homogeneity of the fluorescent background, regular FXm measurements could be performed. We observed that the cell volume changed by a few percent as single cells moved through the collagen gel (**Figure S6G**), with periods of cell protrusion corresponding to a decrease in cell volume. To assess whether these fluctuations in volume were related to the migration of cells, we split individual cells into three groups according to their average speed and plotted their volume (in %) as a function of time (**Figure S6H**). This clearly showed that faster moving cells displayed larger volume fluctuations. Finally, to get a more quantitative assessment of the correlation, we plotted the coefficient of variation of the volume against the speed (**Figure S6I**), for single cells shown in (**Figure S6H**).

Faster cells displayed more volume fluctuations. Interestingly, this relation was well fitted by an extension of the model to cells moving through a meshwork (see **Supplementary file** for the model extension and the fit of the data). This experiment suggests that the mechano-osmotic coupling that we describe in our study is at work in migrating cells, inducing larger volume fluctuations (and thus likely density changes) in faster migrating cells. These volume and density fluctuations could thus be present in a large range of cells in physiological conditions, with yet unknown consequences on cell physiology and behavior.

Beyond the potential functional significance of volume and density fluctuations associated with cell shape changes, our observations and our model demonstrate that a membrane tension homeostasis mechanism is constantly at work in mammalian cells. This mechanism is most likely due to crosstalk between mechanical, osmotic and electrical properties of the cell pointing to the importance of taking into account complex coupling between various physical parameters to understand cellular physics and physiology.

## Acknowledgements

We thank Jian Shi, Rafaële Attia and Li Wang for help with photolithography; Pierre Recho and Romain Rollin for fruitful discussions; Pablo Vargas for help with DCs experiment; Olivier Thouvenin and Matthieu Maurin for assistance with optical details; Vincent Frasier for help with 3D-reconstruction of confined cells. This work was supported by a French Agence Nationale de la Recherche (ANR) grant to MP (ANR-19-CE13-0030). This work has also received the support of Institut Pierre-Gilles de Gennes-IPGG (Equipement d’Excellence, “Investissements d’avenir”, program ANR-10-EQPX-34) and laboratoire d’excellence, “Investissements d’avenir” program ANR-10-IDEX-0001-02 PSL and ANR-10-LABX-31. We also thank IPGG technical platform for providing equipment and technical assistance. LV has received funding from the European Union’s Horizon 2020 research and innovation programme under the Marie Sklodowska-Curie grant agreement no. 641639, and Fondation pour la Recherche Médicale (FDT201805005592). NS acknowledges support from the Human Frontier Science Program (LT000305/2018-L). ASV acknowledges the support of ANR under reference ANR-17-CE13-0020-02.

## Authors contribution

LV. and MP. conceived the study and designed experiments. LV performed the imaging and volume measurements and analyzed the experimental data with help of DC, BD, NS and AD. AS developed the mechano-sensitive PLM model under the supervision of J-FJ and PS and fitted the model predictions to the experimental data. SL performed the tether force measurements under the supervision of AD-M. NS performed mass measurements. AR performed the traction force experiments under the supervision of MB. SV performed electron microscopy experiments. JMGA performed the analysis of cell membranes under confinement. MP supervised the study. LV and MP wrote the manuscript. All authors contributed to the interpretation of the results and commented on the manuscript.**Figure legends**

## Supplementary Material

### Extended Pump and Leak model for cell volume

The volume of the cell changes due to water flux driven by the difference of the osmotic and hydrostatic pressure. The volume dynamics is given by

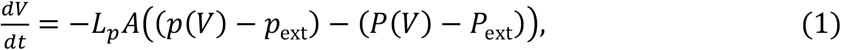

where *L_p_* is the cell s hydraulic conductivity, *A* is the total surface area, *p*_ext_ and *P*_ext_ are the hydrostatic and osmotic pressure of the external medium, respectively. The volume dependence of the hydrostatic *p*(*V*) and osmotic pressure *P*(*V*) of the cell is explicitly shown. The volume dependence of hydrostatic pressure is through the force balance at the cell surface(Cadart et al. 2019) in which both tension and curvature are functions of cell size. In the dilute limit, the osmotic pressure is 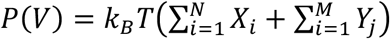, where *T* is the temperature, *X_i_* is the concentration of solute species *i* that are either actively or passively transported across the cell, *Y_j_* is the concentration of impermeant solutes. The concentration of the trapped solute is *Y_j_* = *y_j_*/(*V* – *V*_Solid_) where *y_j_* is the total number of molecules of type j in the cell, and *V*_solid_ is the solid volume of the cell that is inaccessible to the solute molecules. The solid volume is essentially the sum of the volume taken up by the proteins and the DNA. The osmotic pressure of the external medium is 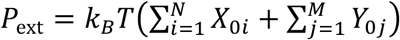.

The hydrostatic pressure in the animal cells is about hundred Pa, whereas the osmotic pressure is three to four orders of magnitude larger. Hence, in **Equation (1)** the hydrostatic pressure can be ignored in comparison to the osmotic pressure. In this limit the volume dynamics is given by

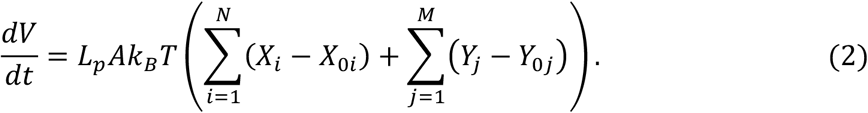

The main solutes that are transported across the cell membrane are sodium, potassium, and chloride(Kay 2017). The sodium and potassium are actively transported through sodium-potassium (Na^+^/K^+^) pumps. The pump transports two potassium into the cell in exchange for three sodium ions transported out of the cell. This leads to the enrichment of potassium inside the cell and sodium outside the cell. Along with the Na^+^/K^+^ pump, there are various cotransporters and channels that passively transport the other ions - chloride, hydrogen ion, carbonates, etc. The ion transport considering only the ion channels and pumps is given by the equation

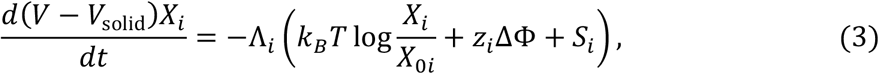

where the first and the second terms are passive flux due to the electrochemical potential difference, ΔΦ is the electric potential energy difference between the inside and the outside of the cell, *z_i_* is the electric charge of species *i*, and *S_i_* is the source term due to active pumping. The volume dependence of the permeability Λ_*i*_ and *S_i_* may be due to the transporters’ mechanosensitivity or due to feedback from signaling molecules that are sensitive to volume change.

The membrane potential is determined using the electroneutrality condition in the cell: 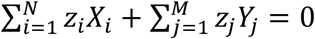. Grouping the positive and negative ions, we get

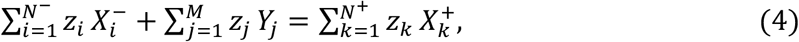

where *N*^−^ is the number of negative ions and *N*^+^ is the number of positive ions. Using the electroneutrality condition and **Equation (3)** the potential is given by

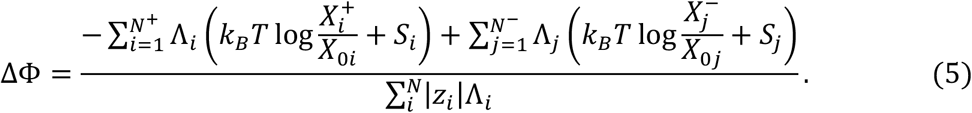

For monovalent ions, i.e., |*z_i_*| = 1, substituting **Equation (4)** in **Equation (2)** we get steady-state volume as

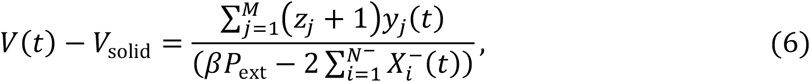

where 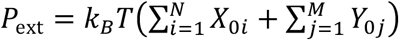, and β = 1/*k_B_T*.

#### Fast timescales: Cell as an osmometer

At the isotonic steady-state condition, the cell volume as given by **Equation (6)** is

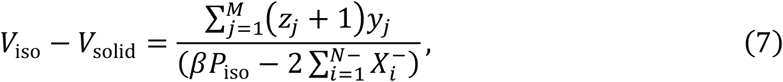

where *P*_ext_ = *P*_iso_ is the osmotic pressure of the external medium at the isotonic condition. For hypotonic shock the medium is diluted leading to osmotic pressure *P*_ext_ = *r*_hypo_*P*_iso_ where *r*_hypo_ is the dilution factor, and for hyperosmotic shock, PEG400 is added to the external medium leading to osmotic pressure *P*_ext_ = *r*_hyper_*P*_iso_, where *r*_hyper_ = (1 + *P*_PEG_)/*P*_iso_. After the osmotic shock, the cell volume changes in response to the new extracellular osmolarity. There is a clear separation in the timescale between water transport and ion transport(Cadart et al. 2019). The ion concentration changes over minutes whereas the volume change due to water flux is on the timescale of seconds. This timescale separation divides the volume dynamics into “fast” passive response, in which the water flows in and out of the cell with constant number of ions within the cell, and “slow” response, in which the ions are transported across the cell. Over the timescale of seconds, the number of ions inside the cells is constant, i.e., *X_i_*(*V* – *V*_Solid_) = *X*_i(iso)_(*V*_iso_ – *V*_solid_). Substituting this in **Equation (2)** we get

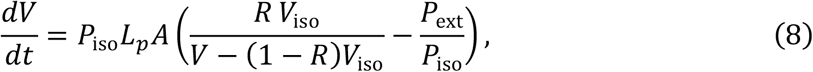

where *R* = (1 – *V*_solid_/*V*_iso_). Thus, we see that the volume dynamics is well approximated by the Van’t-Hoff relation with a fixed number of solutes in the cell. This equation at steady state gives the maximum (minimum) volume (*V_m_*) after the fast hypoosmotic (hyperosmotic) shock. At steady state, we get the Ponder’s relation *V*_m_/*V*_iso_ = *R P*_iso_/*P*_ext_ + (1 – *R*). From **Equation (8)** we get the rate of volume change just after the shock as

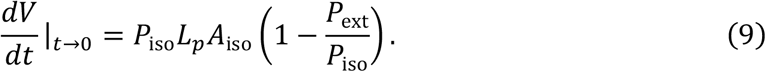

Comparing **Equation (9)** with the experimentally measured rate of volume increase just after the shock, we calculated *L_p_* (**Figure S4C**). The cell volume over the minute’s timescale can be changed from this osmotic shock value by tuning the ion channels and pumps. We first consider the case when the ion transport does not change before and after the osmotic shock. The hypo-osmotic shock in the experiments is attained by dilution, we can see from **Equation (5)** that the membrane potential is constant, and from **Equation (3)** we see that the right had side remains constant. This implies that after the fast increase in volume, the ions reach a new concentration, which is their steady state value, hence, there is no further ion flow at long timescale. However, in the experiments, it has been observed that the cell goes through a volume decrease. This necessarily implies a feedback mechanism for regulatory volume decrease.

In hyperosmotic shock condition, the membrane potential does change; hence, at long times, the volume of the cell increases to a value larger than the maximum decrease. This value is still less than the isotonic volume. The experiments show almost perfect adaptation, implying a regulatory volume increase.

#### Mechano-osmotic mechanism (MOM) for cell volume regulation

There is a clear timescale separation between fluid flow, which is of the order of seconds, and spreading kinetics, which is of the order of minutes. Hence, over the timescale of spreading the cell is in osmotic balance with the external medium. The volume of the cell changes quasi-statically with the change in ion concentration according to **Equation (6)**. The change in volume can be due to change in the number of impermeant ions, or due to a change in the concentration of negatively charged ions. The rate change of the volume, obtained by taking the time derivative of **Equation (6)** is

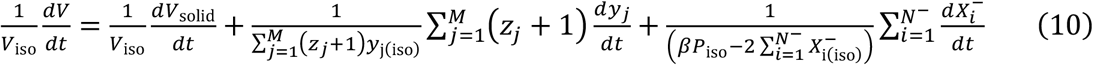

where the first and second term on the right is due to a change in the number of impermeable molecules in the cell due to growth and the third term are due to the change in ion concentration due to change of ion transport rates. The volume-dependent feedback could affect either of the terms. We assume that the change in volume due to feedback is on the ion transport parameters, and the change in impermeable ions is only due to growth. We take the growth rate to be a *r*_growth_ for all the trapped molecules, i.e., *dy_j_*/*dt* = *r*_growth_*y_j_*. The growth in the number of trapped molecules also contributes to the growth in the volume of the solid fraction, for simplicity we take, *dV*_solid_/*dt* = *r*_growth_*V*_solid_. Substituting the growth relations in **Equation (10)** we get constant which gives

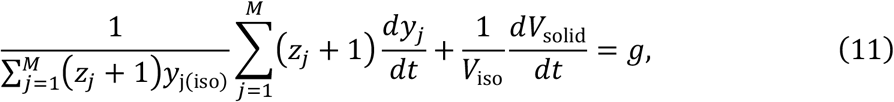

where we have defined the net growth rate of volume as *g* = *r*_growth_ + (1 – *R*) *r*_growth_. In the following we take *g* = 0.05/hr, which gives a doubling time of twenty hours.

Our hypothesis is that the cell shape changes governed by the cytoskeleton rearrangements generate a transient increase in membrane tension; this increase in tension leads to the activation of mechanosensitive ion channels and induces ion flux leading to volume change.

Slow spreading would then induce a lower transient tension increase that would lead to a smaller volume loss. We use a simplified model that includes the mechano-sensitivity of ion transport. Assuming that the ion transport parameters vary quasi-statically, we use the following phenomenological expression for the change in ion concentration:

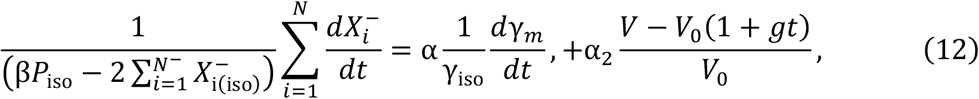

where the term on the right accounts for the change in ion concentration due to the feedback from membrane tension γA on mechanosensitive ion transporters. This defines the mechanosensitivity parameter α. Substituting **Equation (11)** and **Equation (12)** into **Equation (10)** we get

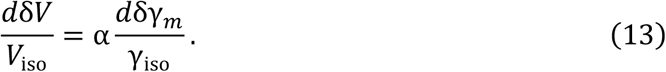

Where *δV* = *V* – *V*_iso_(1 + *g t*) and *δγ_m_* = *γ* – *γ*_iso_. Thus we see that the change in volume is proportional to change in tension. For volume to decrease upon an increase of tension, the coefficient *α* should be negative. For a simplified model of transport of three ions-chloride, sodium, and potassium, we later show that *α*, in general, can take both positive and negative values. For the physiological value of the parameters, we find that *α* be negative if the increase in potassium permeability is much larger than that of sodium.

#### Sign of tension and volume coupling

We now solve for volume, electric potential, and *α*. The external medium is composed of just the ions, sodium, potassium, and chloride. The cytoplasm is composed of these three ions as well an impermeant negatively charged species. Following references(Tosteson and Hoffman 1960; Kay 2017; Kay and Blaustein 2019; Adar and Safran 2020), we consider a simplified model of ion transport. The dynamics of ion transport through the cell membrane as given by **Equation (3)** reads

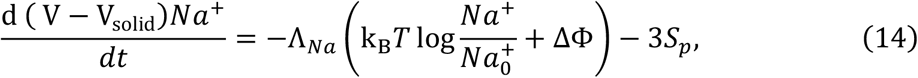

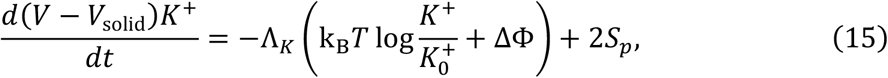

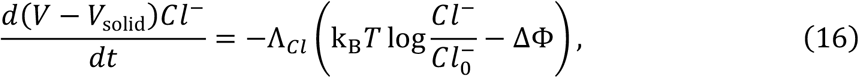

where ΔΦ = Φ – Φ_0_ is the electric potential energy difference between inside and outside of the cell, Λ_*X*_ is the effective permeability of the membrane to ion *X* due to ion channels, and *S_p_* is the activity of the sodium-potassium pump – Na^+^/K^+^ ATPase, the prefactors - minus three and two - are due to the fact that the pump exchanges two potassium for three sodium. Cells pump sodium out of the cell, implying a positive value of *S_p_*. Since the chloride ions are not actively transported there is no active pump contribution to the chloride flux. At steady state, **Equation (14)** to **Equation (16)** gives

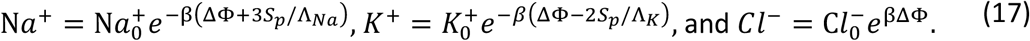

In **Equation (2)** we take *M=1*, the number of impermeable species in the cell is *y*, and *z* is the average negative charge on the molecule. This includes the proteins and the small impermeant charged molecules like phosphate ions. At steady state the osmotic balance reads

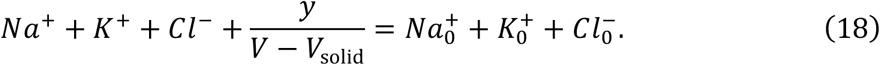

The electroneutrality condition inside the cell and in the external medium is given by *Na*^+^ + *K*^+^ = *Cl*^−^ + *z y*/(*V* – *V*_solid_) and 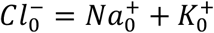, respectively. From the electroneutrality condition, **Equation (17)**, and **Equation (18)** we get the potential difference across the cell membrane, and the cell volume. The volume thus obtained reads

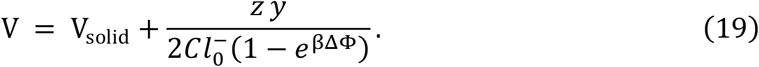

For volume to be finite we need ΔΦ < 0, consistent with different experimental measurements. Substituting **Equation (19)** in the electroneutrality condition inside the cell we obtain a quadratic equation for *e*^βΔΦ^. However, only one of the two roots leads to ΔΦ < 0. The electric potential difference thus obtained reads

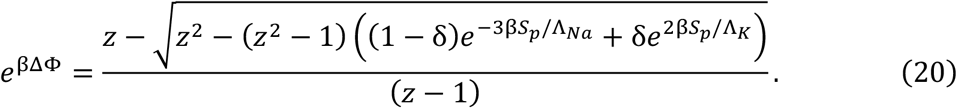

We can now evaluate the change in concentration of chloride ions when tension is changed. Identifying *X_i_*^−^ with *Cl*^−^ the l.h.s of **Equation (12)** reads

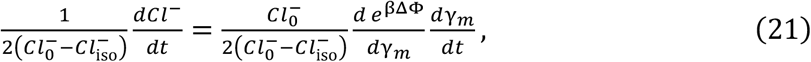

where we have used 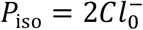. The electric potential difference depends on tension due to mechanosensitivity of the ion channels and pumps. Varying **Equation (20)** we get

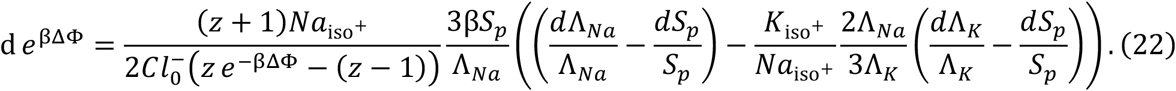

For a small change in tension the channels and pumps change by a small value given by the relation

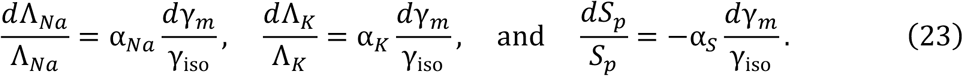

We expect an increase in the channel values and decrease in the pump values due to an increase in tension, implying that the proportionality factors should be positive. Substituting this we get

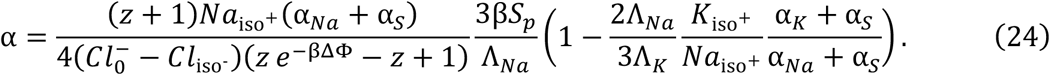

Substituting the parameter values from **Table V** into **Equation (24)** we get

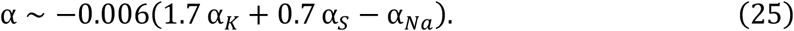

Thus we see that we can get *α* < 0 for comparable values of *α_K_, α_Na_*, and *α_S_*. For slightly different parameters values, it is indeed possible that *α* > 0. Form fitting the experimental volume and spreading data we estimate *ξ* = *A*_0_*kα*/*γ* ~ 1, taking *α* ~ 10^-3^, this gives *A*_0_*k*/*γ* ~ 10. In other words, a 10% increase of contact area leads to a doubling of tension in the elastic regime.

#### Membrane tension as a function of rate change of contact area

We model membrane as a Maxwell viscoelastic element (elastic at short time and viscous at long time), taking the change in membrane tension *γ_m_* to be proportional to the contact area *A_c_*, i.e.,

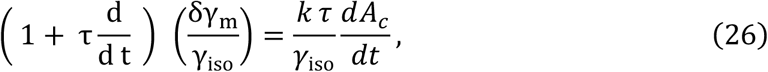

where *k* is the elastic modulus and *τ* is the tension relaxation timescale that depends on various factors related to cortex organization as well on the membrane turnover. Note that *k* is not the elastic response of the lipid bilayer, it is an effective parameter that is related to the cells ability to access its membrane reservoirs upon stretching. Substituting **Equation (26)** in **Equation (13)** we get

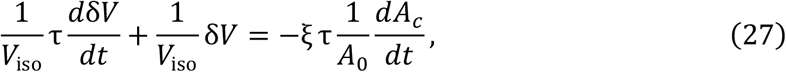

where *ξ* = –*A*_0_*k α*/*γ*_0_. For a time much less than the volume relaxation timescale \tau the rate of volume change is proportional to the rate of spreading

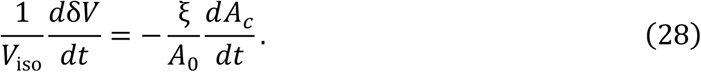

Solving **Equation (27)** we get

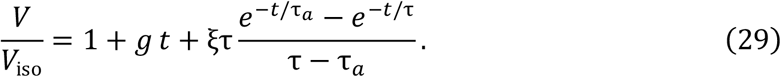

As observed in the experiments, faster the spreading rate larger the volume loss. Since the spreading dynamics are governed mainly by the cortical tension, we can take the time-series of the contact area as input in **Equation (27)**. We first fit the contact area times series to the following equation *A_c_*(*t*) = *A*_0_(1 – *exp*(−*t*/*τ_α_*)), and thus obtain the best fit values of *A*_0_ and *τ_a_*. Substituting this in **Equation (29)** we get volume as a function of time. We then obtain the best fit value of *τ* and *ξ* by numerically fitting the volume dynamics to the measured volume time series using the “Nonlinermodlefitting’’ function in Mathematica.

#### Fast and slow-spreading control cells

As seen from the estimates the drug treatment affects multiple parameters. To confirm the relation between the spreading speed and volume loss we sorted the control cells (n=127) into three equal groups based on their average spreading speed in the first ten minutes. We find that, indeed, the fast-spreading cells lose more volume than the slow-spreading cell. We fit the three groups - slow, intermediate, and fast-spreading cells - to the model and obtain the best-fit parameters. We find that the parameters characterizing the mechano-osmotic feedback (*ξ* and *τ*) are similar for the three classes. Compared to the cells spreading at intermediate speed, the rate of volume loss of the slowest spreading cell was 50% less than that of the fast-spreading cell was 50% more. The average of all the control cells was close to the group of cells with intermediate spreading speed and the fast-spreading cells behaved similarly to the GdCl_3_ treated cells. The value of the parameters is listed in **Table I**.

#### For different drug treatments

The parameter *τ* varies over a wide range of values, whereas *ξ* is about one tenth for all cases except Y-27 and about half for the Y-27 treated cells. For fast-spreading cells treated with Y-27 and GdCl_3_ the spreading rate *A*_0_/*τ_a_* is about 1.8 times that of control. However, the initial rate of volume loss from the model is about four times for Y-27 treated cells and about 1.5 times for the GdCl_3_ treated cells. The parameter *ξ*/*A*_0_ increases by a factor of 1.5 for the Y27 treated cells and decreases by a factor of 0.75 for the GdCl_3_ treated cells. For EIPA treated cells the spreading rate is the same as control but the parameter *ξ* decreases by 20% as seen by a smaller volume loss. The initial rate of volume loss and rate of spreading is similar for the Lat A and CK-666 treated cells.

The volume recovery timescale is quite variable for the difference during treatments. Over the measured timescale of an hour, the volume recovery of the GdCl_3_ and CK-666 treated cells is mainly due to growth. The value of the parameters is listed in **Table II**.

#### Membrane tension as a function of total area

The membrane tension dynamics is given by following Maxwell viscoelastic model:

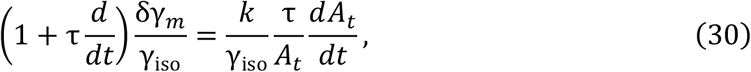

where *γ*_iso_ is the homeostatic value of membrane tension, \tau is the relaxation timescale, *k* is the membrane elasticity, and *A_t_* is the total cell area. Substituting **Equation (13)** in **Equation (30)** the equation for volume dynamics reads

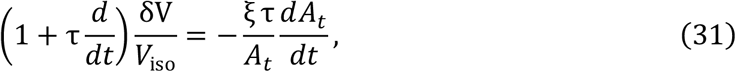

where *ξ* = −*k α*/*γ*_iso_. To compute the total cell area we need to make assumptions about the shape of the cell. At early times, the shape can be well approximated by a spherical cap. However, at a later stage of spreading it is not a reasonable assumption. Since we do not know more about the three dimensional shape of the cell, for simplicity, we model the cell as a spherical cap through the duration of spreading. With this assumption, we can calculate the total area in terms of the instantaneous volume and the contact area. The volume and the total surface area of the spherical cap are given by

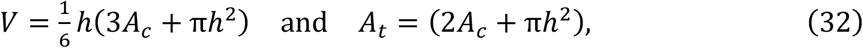

where *h* is the height of the cell and *A_c_* is the contact area. Eliminating the height from **Equation (32)** we get the total area in terms of the volume and contact area, which reads

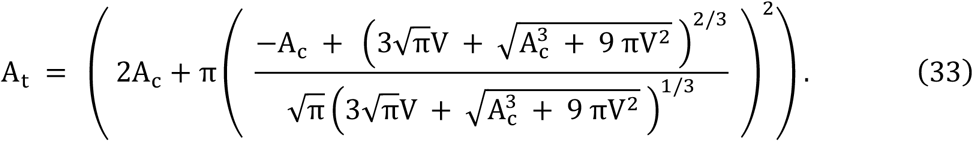

Taking the time derivative of **Equation (33)** and using **Equation (32)** we get the following expression for rate change of the total area as function of rate change of the volume and the contact area:

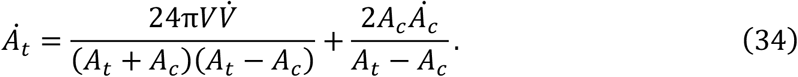

Substituting **Equation (33)** into **Equation (31)** we get

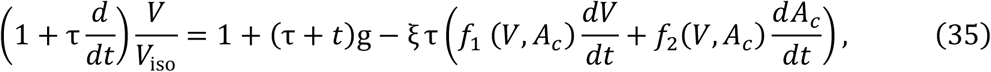

Where 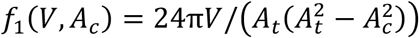 and 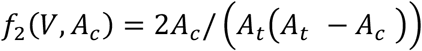.

Rearranging the terms we get

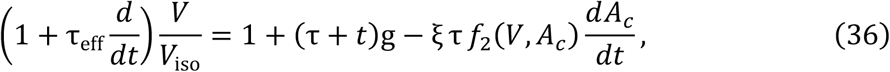

where *τ*_eff_ ≡ *τ*(1 + *ξf*_1_(*V, A_c_*)*V*_iso_). Thus, we see that the volume relaxation timescale is normalized by the volume dependent term in total area, and this effective timescale is larger than the bare tension relaxation timescale.

The effective tension dynamics obtained by by using **Equation (13)** in **Equation (36)** is

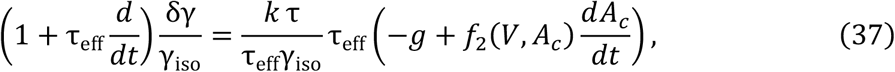

We now fit **Equation (36)** with the experimentally measured volume to obtain the best-fit parameters. For this, we fit the cell spreading data with an exponentially saturating function of the form

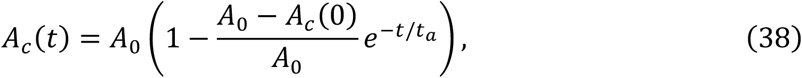

where *A_c_*(0) is equal to the initial contact area which we obtain from the data. We solve **Equation (36)** numerically, using the NDSolve function in Mathematica, for a set of parameter values *ξ* and *τ*. We then select the parameters which minimize the error between the numerically calculated volume and the experimentally measured volume using *L^2^* norm.

The best-fit parameter values for control cells grouped in three groups based on spreading speeds are listed in **Table III** and that for cells treated with different drugs are listed in **Table IV**. We see that the value of *τ* is much smaller compared to the case in which we use contact area as a proxy for total area. This is due to the fact that the volume relaxation time is normalized by the dependence of the total area of the spherical cap on the volume. We also find that parameter *ξ* is an order of magnitude larger compared to the fit using contact area. Although the parameters for the cells in three control groups (Table III) vary significantly, this variation cannot explain the difference in volume loss. If we use the same spreading speed for the three sets of parameters, the resulting volume loss is actually maximum for the parameter corresponding to the slowest spreading cell. This implies that the observed difference in volume loss must be attributed to the difference in spreading speed itself (**Model Figure 1**).

**Model Figure 1.**
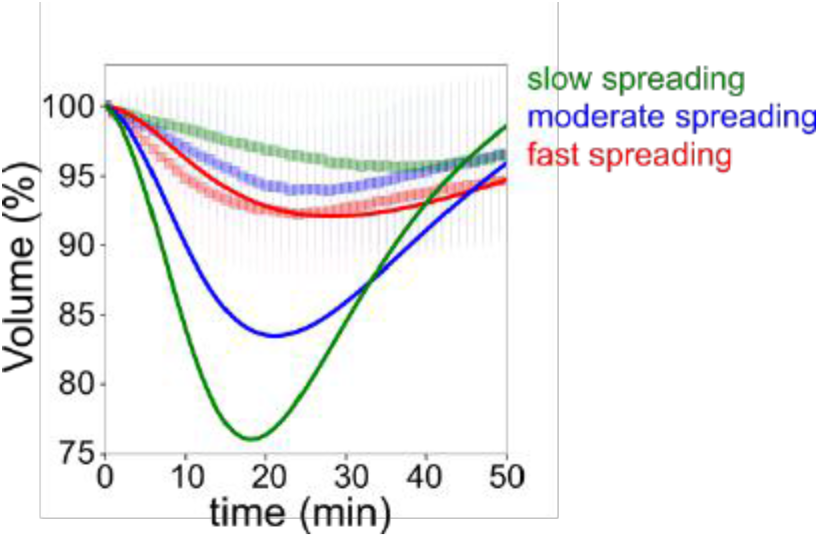
Fits for volume taking the contact area of the fast-spreading cells for the three sets of parameters.

We now compare the change in tension for two cases, one with finite volume tension coupling, i.e, *α* ≠ 0, and compare it with the case *α* = 0, which implies *ξ* = 0. For *ξ* = 0, the volume is given by *V* = *V*_iso_(1 + *g t*).

Substituting this volume in **Equation (30)** we can compute the difference between the two tensions. We find that during the initial spreading the change in tension when there is volume loss due to spreading is always lower than the case when there is not volume loss. In the later part of the spreading, this is not the case. However, since the spherical cap model is more reasonable at the start of the spreading rather than later. This supports the hypothesis that the functional role of the volume loss may be to prevent rapid increase in tension due to fast spreading.

#### Fast Confinement

From **Equation (1)**, we see that, within the PLM framework, for a fixed external medium, the cell volume can change either due to change in hydrostatic or osmotic pressure. The Membrane rupture tension is ~ 20 mN/m, for radius of curvature of about 5 μm this gives Δ*P* ~ 4 × 10^3^ Pa. The change in volume due to this pressure increase as given by **Equation (1)** for constant ion concentration is

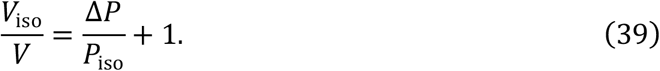

For external osmolarity of 300mM, *P*_iso_ ~ 7 × 10^5^ Pa, which gives Δ*P*/*P*_iso_ ~ 0.5 × 10^-2^o Thus, we see that even at the rupture tension the hydrostatic pressure can change the volume only by about one percent. Hence, the volume loss upon fast confinement that is of the order of ten percent cannot be explained by just the increase of the hydrostatic pressure due to compression.

Within the framework of PLM, the volume change of the order of ten percent can only be due to change in the osmolarity of the cell, which requires transport of ions. For ions transport to take place at timescales of milliseconds, the rates need to increase by four orders of magnitude. Such increase can be easily attained due to pore formation. However, formation of small pores that allow the ions to leak through but does not discriminate between the different ions will lead to an increase rather than a decrease in volume. This is due to the fact that the concentration of ions outside the cell is larger than that inside, hence once the pores open the ion flux is into the cell and the water flux follows the ions flux.

#### Volume fluctuation during cell migration

The cell area fluctuates as the cell passes through the collagen matrix.

We take the total area to be *A_t_* = *A*_0_*sin* (*ωt*), where *ω*~*v_cell_*/*l_mesh_*. As before assuming a viscoelastic model for tension driven by the area *A_t_*, we can compute the volume fluctuation due to mechano-osmotic coupling with membrane tension. The volume dynamics is given by

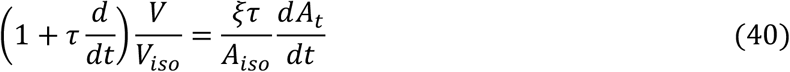

From **Equation (40)** the standard deviation of the volume is

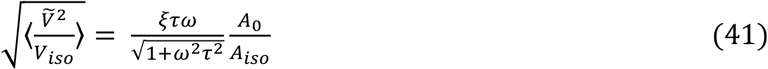

If the distance cell moves in time *τ* is much smaller than the mesh size *v_cell_τ* ≪ *l_mesh_* then the standard deviation of volume increases linearly with the cell velocity. As the speed increases, standard deviation of the volume saturates to the value *ξσ_A_/*A*_iso_*. We fit **Equation (41)** to the experimentally measured values (**Model Figure 2**). Taking the standard deviation of the volume at zero velocity to be 0.01 gives the best fit parameter values to be *τ*/*l_mesh_* = 1.8 *min/μm* and *ξA*_0_/*A*_iso_~0.04. For a mesh size of about five micrometers and area change of the order of few percents we get *τ*~10 *min*, and *ξ*~1, which is in the same range as that obtained when fitting the volume change upon spreading (see **Table IV**).

**Model Figure 2.**
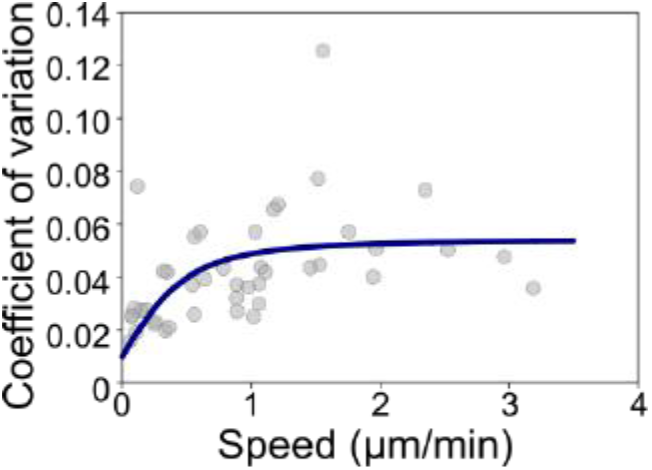
Fits for coefficient of variation of DCs volume using best-fit parameters.

### Model Tables

**TABLE I:** List of fitted parameter values for the control cells are grouped based on spreading speed when the tension depends on change in contact area.

| Condition / Parameters | $A_0 (\mu m^2)$ | $\tau_a (min)$ | $\tau (min)$ | $\xi$ |
| --- | --- | --- | --- | --- |
| Slow spreading | 620 | 63.3 | 98.5 | 0.21 |
| Moderate spreading | 488 | 25.6 | 64.2 | 0.16 |
| Fast spreading | 656 | 17.2 | 99.4 | 0.16 |

**TABLE II:** List of fitted parameter values for different drug treatment when the tension depends on change in contact area.

| Condition / Parameters | $A_0 (\mu m^2)$ | $\tau_a (min)$ | $\tau (min)$ | $\xi$ |
| --- | --- | --- | --- | --- |
| control | 479 | 25.9 | 53.5 | 0.16 |
| Y-27 | 833 | 24.4 | 16.4 | 0.67 |
| EIPA | 549 | 33.7 | 30.7 | 0.18 |
| Lat A | 1522 | 212 | 13.8 | 0.88 |
| GdCl<sub>3</sub> | 628 | 18.3 | 171.2 | 0.17 |
| CK-666 | 285 | 14.9 | ∞ | 0.09 |
| CK-666+Y-27 | 738 | 25.6 | 36 | 0.1 |
| EIPA+Y-27 | 1035 | 29 | ∞ | 0.02 |

**TABLE III:** List of fitted parameter values for the control cells are grouped based on spreading speed when the tension depends on change in total surface area.

| Condition / Parameters | $A_0 (\mu m^2)$ | $\tau_a (min)$ | $\tau (min)$ | $\xi$ |
| --- | --- | --- | --- | --- |
| Slow spreading | 620 | 63.3 | 5 | 2.5 |
| Moderate spreading | 488 | 25.6 | 13 | 0.9 |
| Fast spreading | 656 | 17.2 | 51 | 0.3 |

**TABLE IV:** List of fitted parameter values for different drug treatment when the tension depends on change in total surface area.

| Condition / Parameters | $A_0 (\mu m^2)$ | $\tau_a (min)$ | $\tau (min)$ | $\xi$ |
| --- | --- | --- | --- | --- |
| control | 479 | 25.9 | 12 | 1 |
| Y-27 | 833 | 24.4 | 8.5 | 1.3 |
| EIPA | 549 | 33.7 | 8.5 | 0.8 |
| Lat A | 1522 | 212 | 2 | 3.4 |
| GdCl<sub>3</sub> | 628 | 18.3 | 34.5 | 0.4 |
| CK-666 | 285 | 14.9 | 29.5 | 0.9 |
| CK-666+Y-27 | 738 | 25.6 | 18 | 0.2 |
| EIPA+Y-27 | 1035 | 29 | 4.5 | 0.1 |

**TABLE V:** List of parameter values.

| Parameter | Definition | Estimate |
| --- | --- | --- |
| $Cl_0^-$ | Concentration of chloride ions in the medium | 150 mM(Kay 2017) |
| $K_0^+$ | Concentration of potassium ions in the medium | 3 mM(Kay 2017) |
| $Na_0^+$ | Concentration of sodium ions in the medium | 147 mM(Kay 2017) |
| $Cl_{iso}^-$ | Concentration of chloride ions in the cell | 4 mM (**Equation (17)**) |
| $K_{iso}^+$ | Concentration of potassium ions in the cell | 130 mM (**Equation (17)**) |
| $Na_{iso}^+$ | Concentration of sodium ions in the cell | 5 mM (**Equation (17)**) |
| $Y$ | Concentration of trapped particles in the cell | 30 mM (osmotic balance) |
| $zY$ | Concentration of trapped charges in the cell | 131 mM (electro-neutrality) |
| $\Delta\Phi$ | Potential difference across the plasma membrane | -90 mV (**Equation (5)**) |
| $k_B T$ | Temperature (25°C) | $4.1 \cdot 10^{-21}$ J |
| $\Lambda_{Na}$ | Permeability of $Na^+$ | $2 \cdot 10^{-8}$ moles/(m<sup>2</sup>*s)(Kay 2017) |
| $\Lambda_K$ | Permeability of $K^+$ | $6 \cdot 10^{-7}$ moles/(m<sup>2</sup>*s)(Kay 2017) |
| $\Lambda_{Cl}$ | Permeability of $Cl^-$ | $4 \cdot 10^{-7}$ moles/(m<sup>2</sup>*s)(Kay 2017) |
| $\beta S_p$ | NaK Atpase pumping rate | $1.4 \cdot 10^{-7}$ moles/(m<sup>2</sup>*s) |
| $L_p$ | Hydraulic conductivity | $10^{-(12-13)}$ m/(Pa*s) (**Equation (9)**) |

### Materials and methods

#### Cell culture and drug treatment

HeLa EMBL, HeLa LifeAct, HeLa-MYH9-GFP-LifeAct-mcherry, HeLa Myrpalm-GFP-LiFeact mCherry, Hela hgem-mCherry, RPE-1, 3T3-ATCC cells were maintained in Dulbecco’s Modified Eagle Medium with Glutamax (DMEM/Glutamax; Gibco) supplemented with 10% FBS (Life Technologies) and 1% penicillin-streptomycin solution (Life Technologies), and stored at 37 °C and 5% CO_2_.

Bone marrow derived dendritic cells (DCs) were obtained by differentiation of bone marrow precursors for 10 days in DCs medium (IMDM-Glutamax, FCS 10%, pen-strep 100 U/ml, and 2-ME 50 μM) supplemented with granulocyte-macrophage colony stimulating factor (GM-CSF)-containing supernatant (50 ng/ml) obtained from transfected J558 cell line, as previously described(Barbier et al. 2019).

Latrunculin A (Sigma-Aldrich), CK-666 (Sigma-Aldrich), EIPA (Tocris Bioscience) dissolved in DMSO (Sigma-Aldrich), Y-27632 (Tocris Bioscience), GdCl_3_ (Sigma-Aldrich) dissolved in H_2_O. Incubations with drugs were done for suspended cells 30 min prior experiment, except Latrunculin A added right before experiment to the cells incubated 30 min in medium.

For volume measurements, 10 kDa dextran conjugated with different fluorophores were used in the final concentration 1 mg/ml: fluorescein isothiocyanate–dextran (Sigma-Aldrich), Texas Red (ThermoFisher), Alexa Fluor 647 (ThermoFisher).

For serum starvation experiments, plated cells were incubated overnight in DMEM without FBS. Prior the experiments cells were detached with EDTA and resuspended in the DMEM without FBS collected from cells or in the fresh DMEM supplemented with 10% FBS and incubated for 30 min in suspension.

#### Cell cycle stage detection

The cell cycle state of the cells is indicated by the expression of h-Geminin protein which is expressed by cells from the start of S phase until mitosis(Sakaue-Sawano et al. 2013). To quantify the fluorescence of geminin in the nucleus, firstly a background subtraction is performed on the images using the ImageJ software. An ROI is used to define an area containing the background fluorescence in the image. An average value of the ROI is then subtracted from all the frames. Subsequently, a ROI is drawn to drawn as close to the cell, as possible, and then mean gray value is measured across all the frames.

#### Monitoring of cell volume and contact area while spreading

PDMS-chambers were prepared as described in(Cadart et al. 2017). The typical height of PDMS chambers for volume measurements was 20 μm. PDMS-chambers were incubated with 50 μg/ml fibronectin (Sigma-Aldrich) in PBS for 1 h, washed and incubated overnight with culture medium. Cells were detached with warm Versen (Gibco) and resuspended in medium collected from cells to facilitate spreading.

In case of measurements of non-adherent cells, we used chambers incubated with PLL-PEG coating (0.1 mg/ml solution in HEPES, SuSoS), washed and incubated overnight with culture medium without FBS. Cells were detached with Trypsin and resuspended in a fresh culture medium.

The cell volume measurement explained in details in(Cadart et al. 2017) and used in were coupled with spreading area measurement performed by Reflection Interference Reflection Microscopy (IRM)(Rädler and Sackmann 1993; Cuvelier et al. 2007). Microscopy was performed at 37 °C with 5% CO_2_ atmosphere. Imaging was started immediately after cell injection into the chamber with 1 min time interval. Imaging was performed using a ZEISS Z1 Observer epi-fluorescence microscope equipped with an Orca-Flash 4 Camera (Hamamatsu), 20X Plan-Apochromat objective, NA0.8 and the software Metamorph (Molecular Device).

The volume extraction was performed with a MatLab software as described in(Cadart et al. 2017).

The analysis of spreading and contact area was performed manually using the ImageJ software. The borders of the cell were delimited manually and then the area, and different shape descriptors were extracted.

For HeLa and 3T3 cells initial speed of spreading 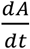 and volume flux 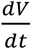 was calculated as linear slope in the first 10 min after measurable cell to substrate contact. For RPE-1 cells initial 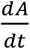 and 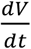 were calculated as linear slopes in the 10 min (or less, if it happen early) prior time point when spreading area is equal to cross-section area of cell in initial non-spread state, and in the first 10 min after that time point.

#### Micropatterning

Cells were patterned using the existed technique(Azioune et al. 2011).

#### Side-view microscopy

Glass slide was attached to glass bottom dish by UV-glue, the position of glass was slightly tilted from perpendicular to the dish bottom. Glass was coated with fibronectin and washed with medium. Cells were detached with Versen and resuspended in warm medium collected from cells and incubated 30 min. Then drop of cell was added to the dish, close to the angle between dish bottom and attached glass. Dish was placed to the incubator for 2 minutes to allow cell initial attachment to the tilted glass. Then 2 ml of medium collected from cells were added to the dish and microscopy started with time frame 1 min. Imaging was performed using a ZEISS Z1 Observer epi-fluorescence microscope 20X NA0.4.

#### Monitoring of cell volume during cell migration in the collagen

Collagen mix was prepared on ice to delay polymerization: 25 μl 10X PBS + 25 μl culture medium + 55 μl collagen + 140 μl culture medium with DCs (2*10^6^/ml) + 5 μl FITC-dextran + 1.3 μl NaOH

Immediately after mixing suspension was added into PDMS-chamber for volume measurements with height 12 μm. Microscopy was started ~10 min after injection. Imaging was performed using a ZEISS Z1 Observer epi-fluorescence microscope.

Cell velocity during migration in collagen gel was calculated for 10 min intervals. Cell position was defined as a center of mass of a binary mask applied on FXm images of cells.

#### Monitoring of cell volume during osmotic shock

PDMS-chambers were coated with 0.01% PLL (Sigma-Aldrich) to prevented cell detachment during changing medium and maintaining cell round shape during experiment, then washed and incubated overnight with culture medium without FBS. Cells were detached with Trypsin. Isoosmotic medium was exchanged to the medium with known osmolarity typically 2.5 min after beginning of acquisition. Full medium exchange in the chamber takes less than 1 s. Imaging was performed using a ZEISS Z1 Observer epi-fluorescence microscope equipped with 20X NA0.4. Hypoosmotic solutions were made by water addition to culture medium, hyperosmotic by addition PEG400. Osmolarity of working solutions was measured by osmometer Type 15M (Löser Messtechnik).

Cell rupture in response to distilled water exposure was monitored by propidium iodide (1 μg/ml) (Sigma-Aldrich) intensity inside the cell.

Volume flux for passive response to osmotic shock was defined as a linear slope at the linear region of volume curves.

Adaptation speed for osmotic shock recovery was calculated as a linear slope at 5 min intervals.

#### Monitoring of cell volume under confinement

Cells were detached with Trypsin, resuspended in fresh culture medium. Both static 6-well confiner and dynamic confiner were used according to experimental procedure described in (Le Berre et al. 2014). Imaging was performed using a ZEISS Z1 equipped with 20X Long-Distance objective NA0.4.

For volume measurements performed with dynamic confiner, bottom glass was coated with 0.01% PLL that prevented cell escape from the field of view and allowed following the same cells before and after confinement.

Calculation of surface area of non-confined cell were done with the assumption of spherical cell shape, and of confined cells with the assumption of cylindrical cell shape, based on measured cell volume.

#### Spinning disk microscopy

Qualitative imaging for osmotic shock and confinement experiments was performed with spinning disk set-up (Leica DMi8). 63X and 100X oil objectives were used. CellMask (Invitrogen) staining was performed in warm PBS solution (1 μl of dye to 1000 μl PBS).

Filopodia were manually segmented. Filopodia density is plotted as number of filopodia per μm of cell body diameter. Bleb were manually segmented from middle plane images. For membrane density measurements on cell contour cells were background substrated, and resliced by their contour, where most of the membrane marker accumulates. An average projection was plotted for 3μm around the cell edge.

#### Dry mass measurements

Mass measurement was performed by quantitative phase microscopy using Phasics camera(Aknoun et al. 2015). Images were acquired by Phasics camera every 15 min for 35 hours during the duration of the experiment. To get the reference image, 32 empty fields were acquired on the PDMS chips and a median image was calculated. Custom MATLAB scripts were written by Quantacell for analysis of interferograms (images acquired by phasics). The interferograms were associated with reference images to measure the optical path difference and then separated into phase, intensity and phase cleaned images (background set to 1000 and field is cropped to remove edges). Background was then cleaned using gridfit method and a watershed algorithm was used to separate cells which touch each other. Mass was then calculated by integrating the intensity of the whole cell.

#### Tether pulling

For apparent membrane tension measurements, tether force was measured with single cell atomic force spectroscopy by extruding tethers from the plasma membrane of HeLa Kyoto cells. Cellview^®^ glass bottom dishes (Greiner) were coated for 1 h with fibronectin (50 μg/ml; Sigma-Aldrich). Cells were then plated in the presence of drugs or vehicle, and probed either during spreading (30 min after plating) or at steady state (fully spread; 4 h after plating).

Tether extrusion was performed on a CellHesion^®^ 200 BioAFM (Bruker) integrated into an Eclipse Ti^®^ inverted light microscope (Nikon). OBL-10 Cantilevers (spring constant ~60 pN/nm; Bruker) were mounted on the spectrometer, calibrated using the thermal noise method (reviewed in(Houk et al. 2012)) and coated with 2.5 mg/ml Concanavalin A (Sigma-Aldrich) for 1 h at 37° C. Before the measurements, cantilevers were rinsed in PBS and cells were washed and probed in Dulbecco’s Modified Eagle Medium with Glutamax (DMEM/Glutamax; Gibco) supplemented with 2% FBS (Life Technologies) and 1% penicillin-streptomycin solution (Life Technologies). Measurements were run at 37° C with 5% CO_2_ and samples were used no longer than 1 h for data acquisition.

Tether force was measured at 0 velocity at which is linearly proportional to apparent membrane tension, assuming constant membrane bending rigidity(Hochmuth et al. 1996). In brief, approach velocity was set to 0.5 μm/s while contact force and contact time ranged between 100 to 200 pN and 100 ms to 10 s respectively, aiming to maximize the probability to extrude single tethers. To ensure tether force measurement at 0 velocity, the cantilever was retracted for 10 μm at a velocity of 10 μm/s. The position was then kept constant for 30 s and tether force was recorded at the moment of tether breakage at a sampling rate of 2000 Hz. Resulting force-time curves were analyzed using the JPK Data Processing Software.

#### Traction force measurements

Force measurements were conducted directly after seeding the cells on the sample and spreading was observed for 90 minutes on an inverted microscope (Nikon Ti-E2) with a Orca Flash 4.0 sCMOS camera (Hamamatsu) and a temperature control system set at 37°C. To avoid shaking the cells during stage movement, a POC-R2 sample holder in closed perfusion configuration was used and cells were seeded with a syringe right before image acquisition. The medium was supplemented with 20 mM of HEPES in order to buffer the pH during the experiment. Force measurements were performed using a method described previously(Tseng et al. 2011). In short, Fluorescent beads were embedded in a polyacrylamide substrate with 20 kPa rigidity and images of those beads were taken during cell spreading. The first frame, before cells started attaching to the substrate, served as unstressed reference image. The displacement field analysis was done using a homemade algorithm based on the combination of particle image velocimetry and single-particle tracking. After correcting for experimental drift, bead images were divided into smaller subimages of 20.7 μm width. By cross correlating the sub-images of the stressed and the unstressed state, mean displacement of the sub-image can be measured. After correcting for this displacement, the window size is divided by 2 and the procedure is repeated twice. On the final subimages, single-particle tracking was performed to obtain a subpixel resolution displacement measurement. From the bead displacement measurements a displacement field was then interpolated on a regular grid with 1.3 μm spacing. Cellular traction forces were calculated using Fourier transform traction cytometry with zero-order regularization(Sabass et al. 2008; Milloud et al. 2017), under the assumption that the substrate is a linear elastic half-space and considering only displacement and stress tangential to the substrate. To calculate the strain energy stored in the substrate, stress and displacement field were multiplied with each other and with the grid pixel area and then summed up over the whole cell. All calculations and image processing were performed with MATLAB.

#### Electron microscopy

Hela cells plated on fibronectin-coated glass coverslips for 30 minutes were disrupted by scanning the coverslip with rapid sonicator pulses in KHMgE buffer (70 mM KCl, 30 mM HEPES, 5 mM MgCl2, 3 mM EGTA, pH 7.2). Paraformaldehyde 2%/glutaraldehyde 2%-fixed cells were further sequentially treated with 0.5% OsO4, 1% tannic acid and 1% uranyl acetate prior to graded ethanol dehydration and Hexamethyldisilazane substitution (HMDS, Sigma-Aldrich). Dried samples were then rotary-shadowed with 2 nm of platinum and 5-8 nm of carbon using an ACE600 high vacuum metal coater (Leica Microsystems). Platinum replicas were floated off the glass by 5% hydrofluoric acid, washed several times by floatation on distilled water, and picked up on 200 mesh formvar/carbon-coated EM grids. The grids were mounted in a eucentric side-entry goniometer stage of a transmission electron microscope operated at 80 kV (Philips, model CM120) and images were recorded with a Morada digital camera (Olympus). Images were processed in Adobe Photoshop to adjust brightness and contrast and presented in inverted contrast.

### Supplementary figures legend

**Figure S1.**
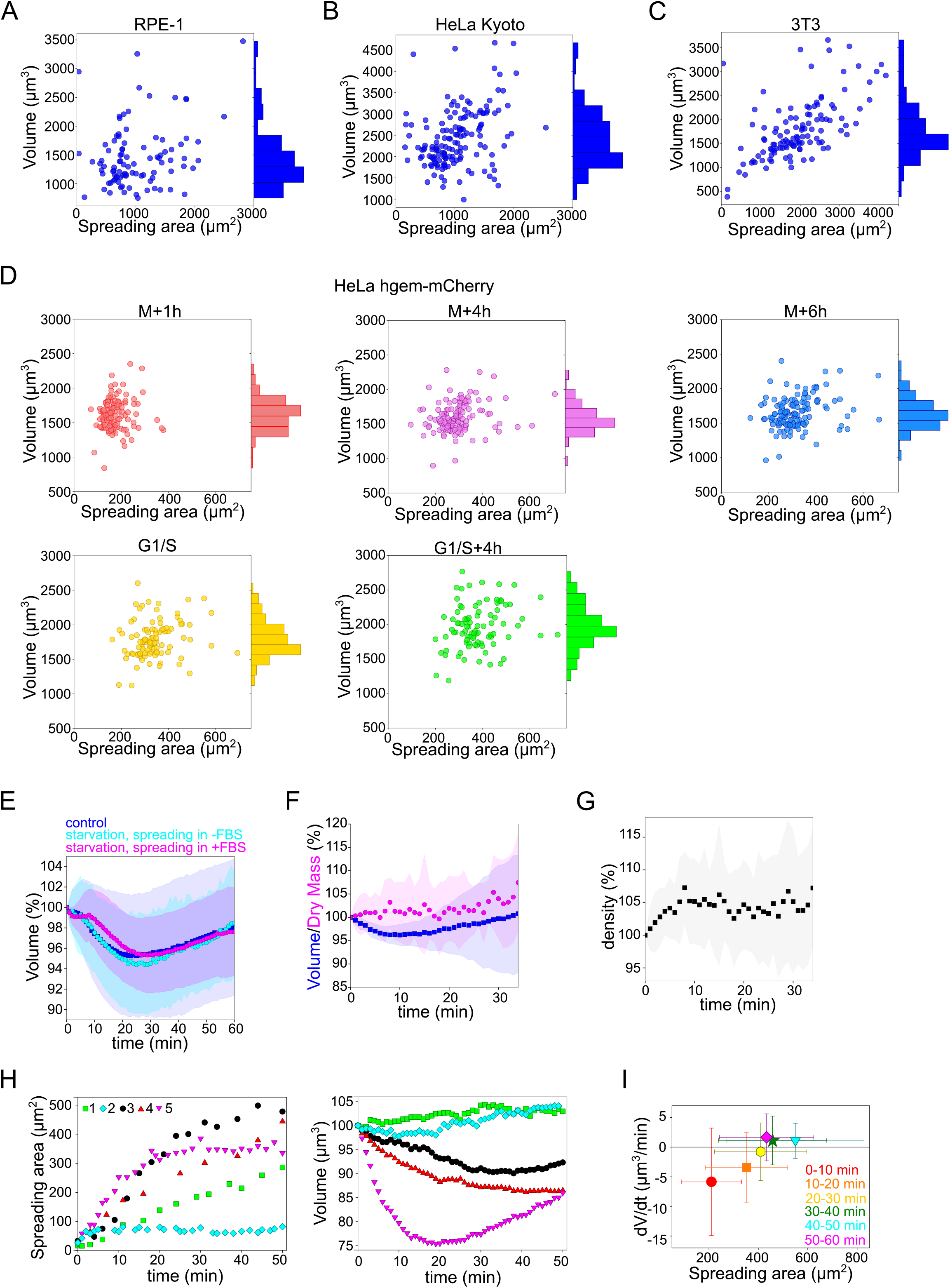
**A.** Relation between volume and spreading area of RPE-1 cells 4h after plating on the fibronectin-coated glass (n=95). R=0.27. **B.**Relation between volume and spreading area of HeLa Kyoto cells 4h after plating on the fibronectin-coated glass (n=151). R=0.35. **C.** Relation between volume and spreading area of 3T3 cells 4h after plating on the fibronectin-coated glass (n=117). R=0.61. **D.** Relation between volume and spreading area of HeLa hgem-mCherry cells at the different cell cycle stages: M+1h (n=131) R=0.11, M+4h (n=131) R=0.23, M+6h (n=131) R=0.26, G1/S (n=99) R=0.20, G1/S+4h (n=92) R=0.22, represented on **Figure 1A**. **E.** Average normalized volume of control HeLa Kyoto cells (blue, n=99), cells incubated overnight in the serum-free medium (cyan, n=61) or cells incubated overnight in the serum-free medium and resuspended in FBS containing medium prior experiments (magenta, n=71) spreading on fibronectin-coated glass. Error bars represent standard deviation. **F.** Average normalized cell volume (blue) and dry mass (magenta) of HeLa Kyoto cells spreading on fibronectin-coated glass (n=17). Error bars represent standard deviation. **G.** Average normalized density of cells represented on the panel **Figure 1SE**. **H. Left:** Spreading area of individual HeLa Kyoto cells during spreading on fibronectin-coated glass. **Right:** Normalized volume of individual HeLa Kyoto cells spreading on fibronectin-coated glass. **I.** Volume flux (dV/dt) of single control HeLa Kyoto cells (n=195) plotted versus their spreading area at the 50-60 min of spreading. Error bars represent standard deviation. Color bar indicate kernel density.

**Figure S2.**
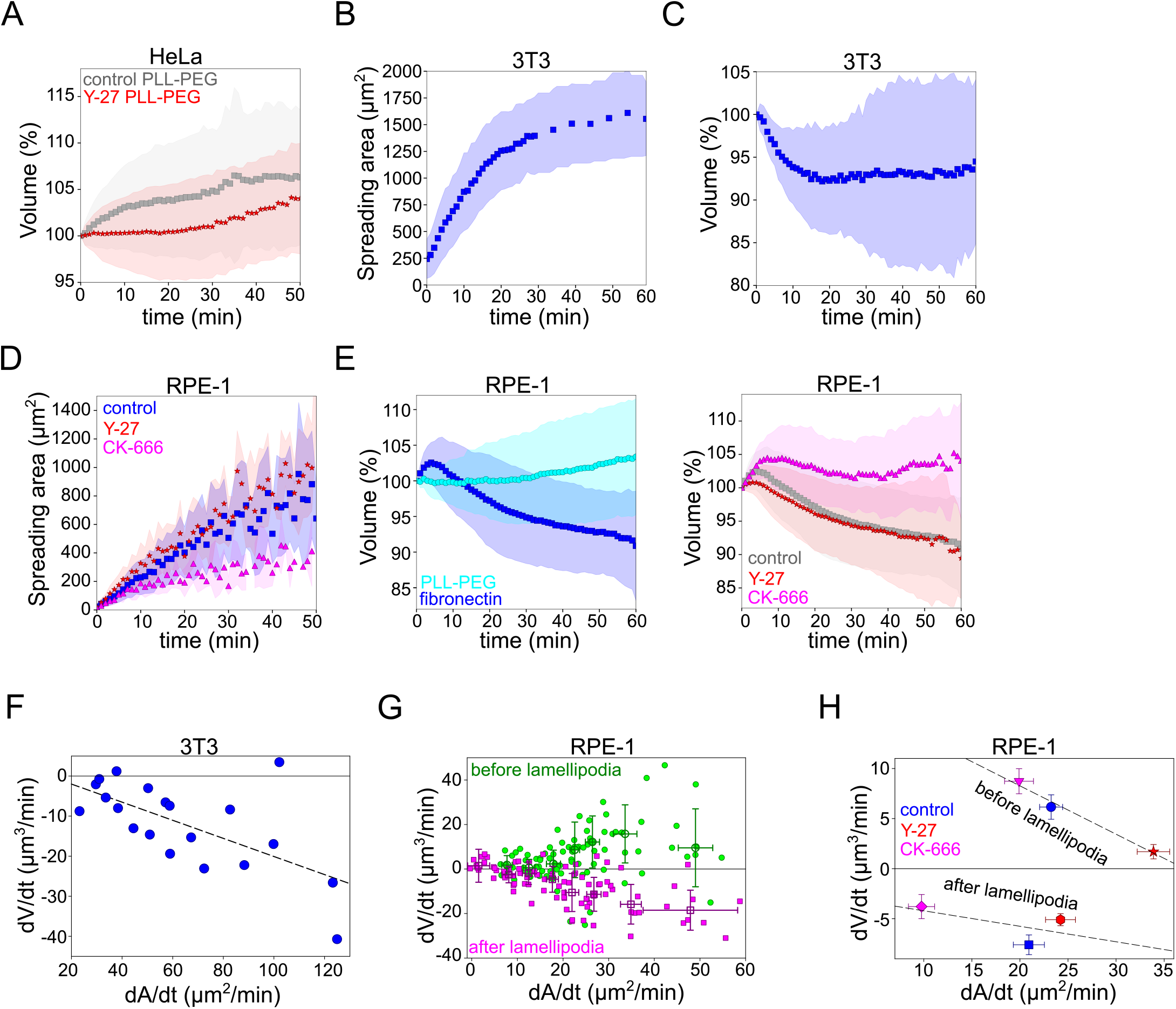
**A.** Average normalized volume of control HeLa Kyoto cells (grey, n=84) or treated with 100 μM Y-27632 (red, n=115) plated on PLL-PEG-coated glass. Error bars represent standard deviation. **B.** Average spreading area of 3T3-ATCC cells spreading on fibronectin-coated glass (n=20). Error bars represent standard deviation. **C.** Average normalized volume of 3T3-ATCC cells spreading on fibronectin-coated glass (n=20). Error bars represent standard deviation. **D.** Average normalized spreading area of control RPE-1 cells (grey, n=90), treated with 100 μM Y-27632 (red, n=85) or 100 μM CK-666 treated (magenta, n=37) spreading on fibronectin-coated glass. Error bars represent standard deviation. **E. Left:** Average normalized volume of control RPE-1 cells (blue, n=90) or control cells plated on PLL-PEG-coated glass (cyan, n=47). Error bars represent standard deviation. **Right:** Average normalized volume of control RPE-1 cells (grey, n=90), 100 μM Y-27632 treated (red, n=85) or 100 μM CK-666 treated (magenta, n=37) spreading on fibronectin-coated glass. Error bars represent standard deviation. **F.** Volume flux (dV/dt) of 3T3-ATCC cells plotted versus their spreading speed (dA/dt) at the first 10 min of spreading (n=20). **G.** Volume flux (dV/dt) of RPE-1 cells plotted versus their spreading speed (dA/dt) before (green) and after (magenta) lamellipodia formation (n=90). Error bars represent standard error. **H.** Median volume flux (dV/dt) of control (blue, n=89), 100 μM Y-27632 (red, n=85) or 100 μM CK-666 (magenta, n=38) treated RPE-1 cells plotted versus their spreading speed (dA/dt) before and after lamellipodia formation. Error bars represent standard error.

**Figure S3.**
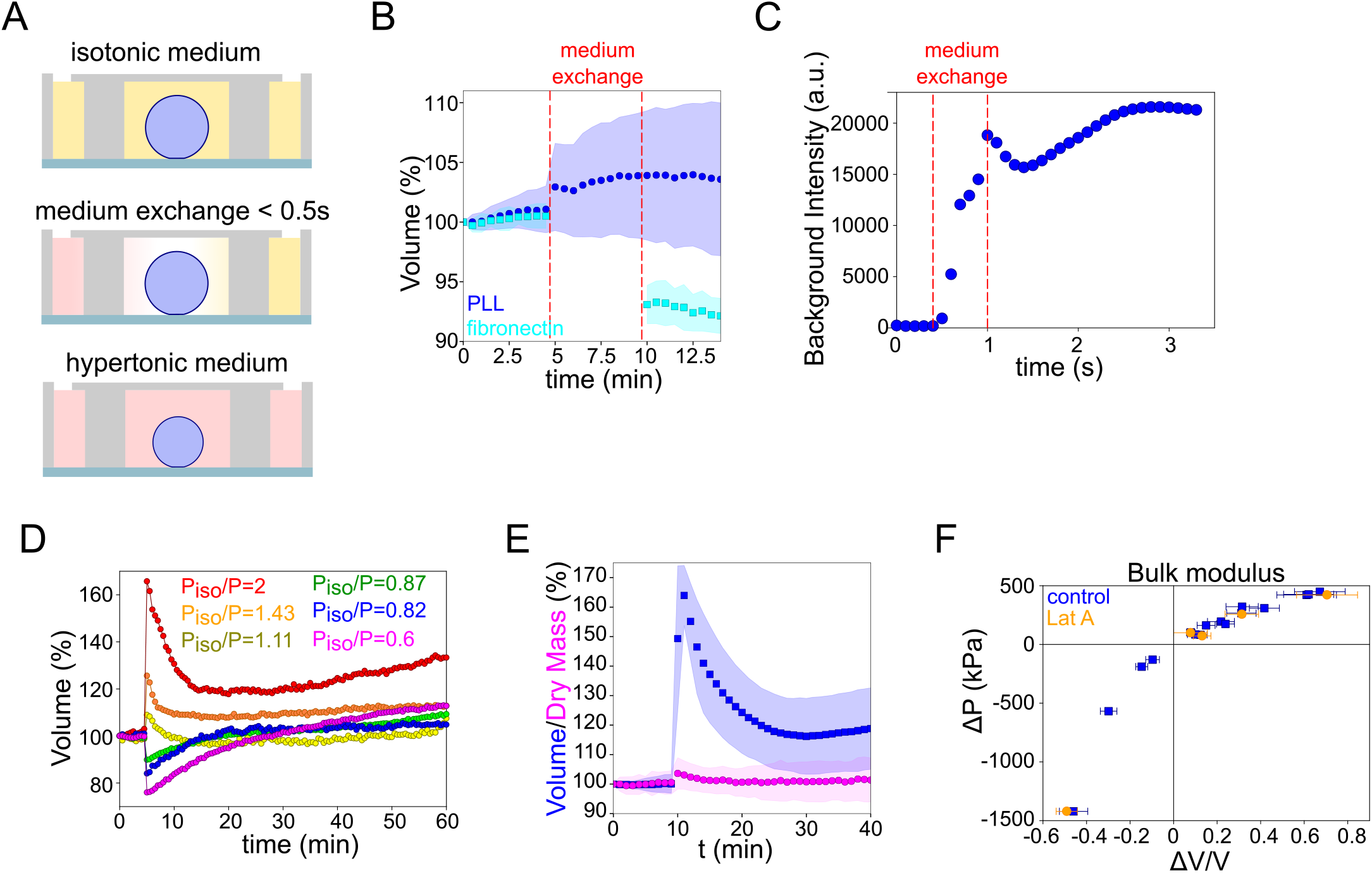
**A**. Schematic of osmotic shock experiments in FXm chamber. **B**. Comparison of cell volume changes of HeLa Kyoto cells in response to the change of medium to the medium with the same osmolarity for the cells adherent on fibronectin-coated (cyan, n=10) and cells attached on PLL-coated (blue, n=35) glasses. Red lines indicate the time of medium exchange. Error bars represent standard deviation. **C**. Timescale of medium exchange inside the volume measurement chamber. Full replace of medium (between 2 dashed lines) inside the chamber occurred in ~0.5 s. **D**. Examples of single HeLa Kyoto cells volume changes in response to osmotic shocks of different magnitudes. **E**. Average cell volume (blue) and dry mass (magenta) of HeLa Kyoto cells in response to hypoosmotic shock (n=63). Error bars represent standard deviation. **F**. Relative changes in osmotic pressure induced by osmotic shock plotted versus deformation for control (blue) and 2 μM Lat A treated (orange) HeLa Kyoto cells. Based on the same data as in **Figure 3E**. Error bars represent standard deviation.

**Figure S4.**
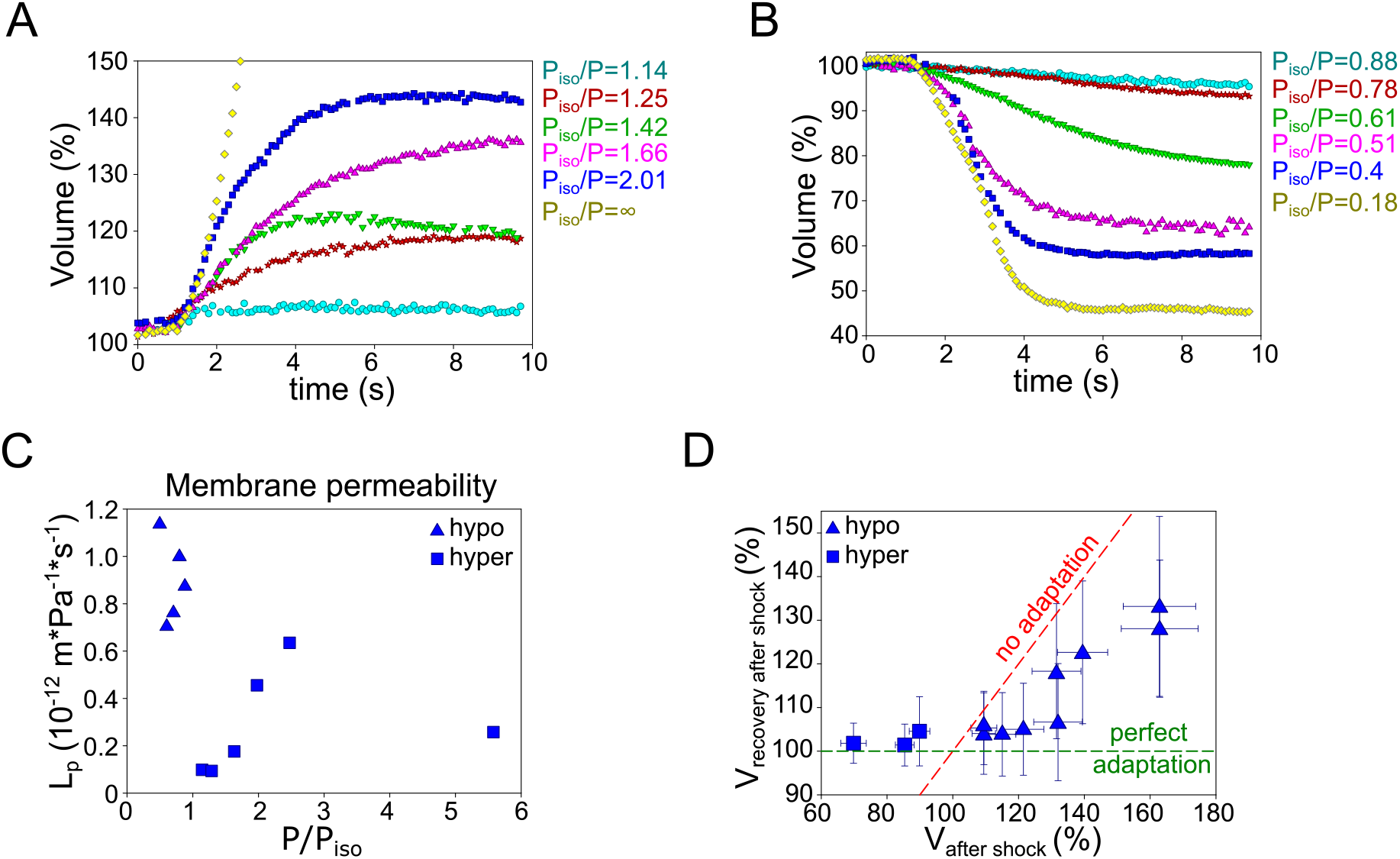
**A.** Volume of single HeLa Kyoto cells during initial response to hypoosmotic shocks of different magnitudes measured with high time resolution. **B.** Volume of single HeLa Kyoto cells during initial response to hyperosmotic shocks of different magnitudes measured with high time resolution. **C.** Estimated hydraulic conductivity for osmotic shock of different magnitudes. Calculations are based on the data presented on **Figure 4B.** **D**. Relative volume changes in HeLa Kyoto cells followed 30 min by passive volume response to osmotic shock plotted versus the values reached during passive response. Green dashed line corresponds to the “perfect adaptation”, red dashed lines correspond to the absence of adaptation. Error bars represent standard deviation.

**Figure S5.**
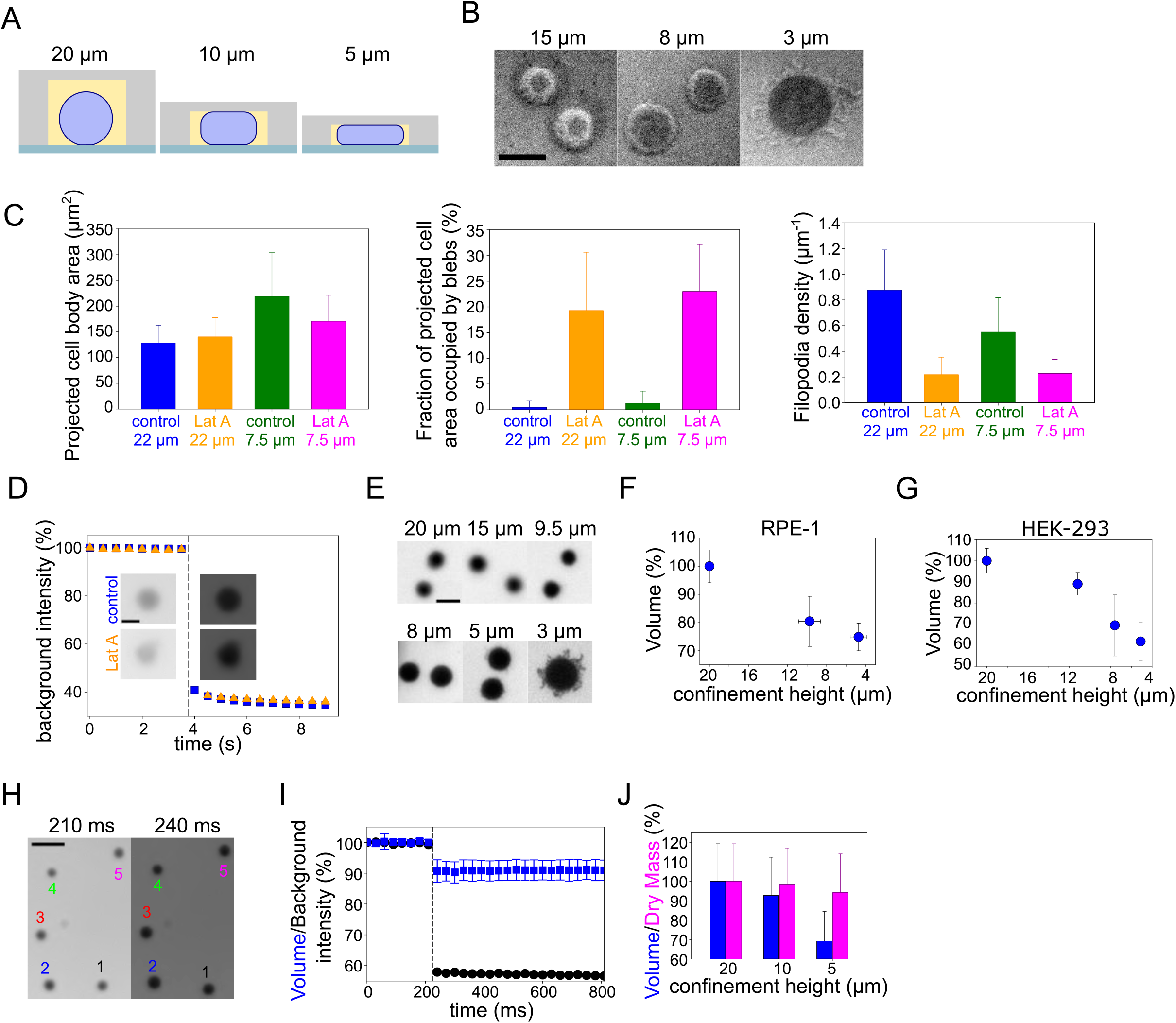
**A.** Schematic of FXM experiments under confinement. **B**. RICM images of cell contact area under different confinement heights. Scale bar 10 μm. **C**. Analysis of control and 2 μM Latrunculin A treated cell morphological changes under confinement. **Left:** Average projected cell body area of cells. **Middle:** Average fraction of projected cell area occupied by blebs. **Right:** Filopodia density. For each panel analysis is performed on the middle Z-plane of HeLa-MYH9-GFP-LifeAct-mcherry cells. Number of cells in each condition n=10. Error bars represent standard deviation. **D**. Corresponding to **Figure 5F** background intensity values and FXm images of single cells before and after confinement. Scale bar 10 μm. Error bars represent standard deviation. **E**. FXm images of Hela Kyoto cells under different confinement heights. Scale bar 10 μm. **F**. Average normalized volume of RPE-1 cells under different confinement heights. Each data point represents an average of N=3 experiments, each experiment contains n~140 individual cells. Error bars represent standard deviation. **G.** Average normalized volume of HEK-293 cells under different confinement heights. Each data point represents an average of N=5 experiments, each experiment contains n~118 individual cells. Error bars represent standard deviation. **H**. Corresponding to **Figure 5I** FXm images. **I.** Corresponding to **Figure 5I** background intensity values (black) and average normalized volume (blue, n=19). Dashed line indicates the moment of confinement. **J**. Average cell volume (blue) and dry mass (magenta) of HeLa Kyoto cells under different confinement heights. Number of single cells analyzed for each height: 20 μm (n=37), 10 μm (n=174), 5 μm (n=64).

**Figure S6.**
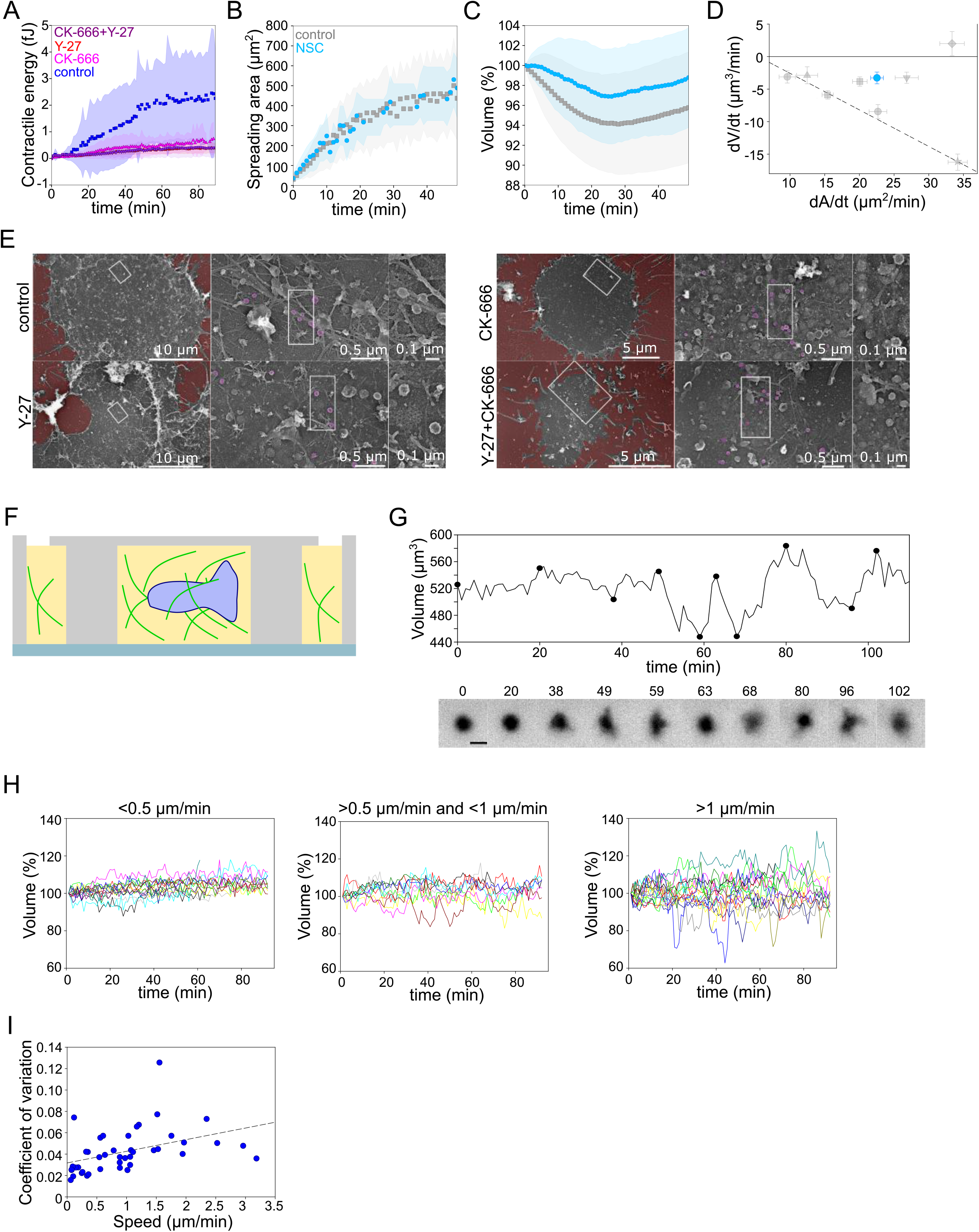
**A.** Contractile energy of control, treated with Y-27, CK-666 or CK-666+Y-27 HeLa EMBL cells during spreading on fibronectin-coated glass. **B.** Average spreading area of control HeLa Kyoto cells (grey, n=125) or 20 μM NSC (blue, n=67). Error bars represent standard error. **C.** Average normalized volume of control HeLa Kyoto cells (grey, n=125), or 20 μM NSC (blue, n=67). Error bars represent standard error. **D.** Volume flux (dV/dt) plotted versus their spreading speed (dA/dt) of single control HeLa Kyoto cells treated with various drugs and represented at **Figure 6K** and treated with 20 μM NSC (n=101). Error bars represent standard error. **E.** Platinum replica electron microscopy survey views of the cytoplasmic surface in control, Y-27632, CK-666 or CK-666 + Y-27632-treated unroofed Hela cells spread on glass coverslips for 30 minutes. Extracellular substrate is pseudo-colored in red. For each panel, high magnification views corresponding to the boxed regions are shown on the right. **F.** Schematic of volume measurements of DCs migrating in collagen in FXm chamber. **G. Top:** Volume of single DC migrating in collagen. **Bottom:** Corresponding FXm images. **H.** Volume of single DCs migrating in collagen with the different speeds. **Left:** < 0.5 μm/min (n=14), **middle:** >0.5 μm/min (n=10) and < 1 μm/min, **right:** > 1 μm/min (n=19). **I.** Coefficient of variation of volume flux dV/dt computed for 10 min intervals during single DCs migration in collagen plotted versus their average speed (n=43).

### Supplementary movies legend

**<u>Movie S1</u>**

Side view of a HeLa-LifeAct cell spreading on fibronectin-coated glass. Top: bright field, bottom: LifeAct. Scale bar 10 μm. 20xPA.

**<u>Movie S2</u>**

Spreading of HeLa EMBL cells on fibronectin-coated glass. Left: FXm, right: RICM. 20xPA.

**<u>Movie S3</u>**

FXm imaging of HeLa EMBL cells attached on PLL-coated glass exposed to osmotic shock. 20xLD.

**<u>Movie S4</u>**

3D-shape reconstruction by FXm of HeLa EMBL cells spread for 20 min at fibronectin-coated glass. Control, Y-27632, CK-666, CK-666+Y-27632. 63x.

**<u>Movie S5</u>**

FXm imaging of HeLa EMBL cells attached on PLL-coated glass exposed to osmotic shock recorded with high frame rate. 20xLD.

**<u>Movie S6</u>**

3D-membrane reconstruction of HeLa expressing MyrPalm-GFP (black) cells cell shape under different confinement heights. 63x.

**<u>Movie S7</u>**

Z-planes of control and 2 μM Lat A treated HeLa-MYH9-GFP-LifeAct-mcherry, cells under 20 μm and 7.6 μm confinement heights. Cell membrane is stained with CellMask Far Red (white). 63x.

**<u>Movie S8</u>**

FXm imaging of HeLa EMBL cells during dynamic confinement recorded with 20xLD.

**<u>Movie S9</u>**

FXm imaging of HeLa EMBL cells during dynamic confinement recorded with 20xPA.

**<u>Movie S10</u>**

FXm imaging of DCs migrating in collagen gel. 20xLD.

